# Combination-Based Drug Screening for Induced Oligodendrocyte Differentiation Enables Mechanistic Insight and Identifies Optimal Drug Pairs for Remyelination

**DOI:** 10.1101/2023.07.11.548469

**Authors:** Brittney A. Beyer, Amanda Sul, Jared T. Gillen Miller, Björn Neumann, Warren C. Plaisted, Toru Kondo, Robin J.M. Franklin, Luke L. Lairson

## Abstract

Remyelination-promoting agents have significant potential utility as therapies for the treatment of demyelinating diseases, including multiple sclerosis. Clemastine and bexarotene have recently been evaluated in Phase II clinical trials to evaluate their potential in this context, with evidence for drug-induced remyelination being observed in both trials. Efficacy levels for both agents as monotherapies, as well as dose-limiting toxicities, highlight the need for more effective approaches. Additionally, questions about the relevance of M1R as the target of clemastine, and also around a mechanism involving accumulation of 8,9-unsaturated sterols, remain. Here, we have identified potent alternatives to clemastine (i.e., doxepin and orphenadrine), which are predicted to have superior tolerability and efficacy profiles and provide mechanistic insight related to M1R, and have completed pairwise drug combination screens using diverse classes of OPC differentiation-inducing agents. Vitamin D receptor agonists were found to enhance M1R antagonist-induced OL differentiation. Select compounds implicated in 8,9-unsaturated sterol accumulation synergistically enhanced the activity of bexarotene in OPCs, which resulted in insights that implicate a critical role for liver-X-receptor in the mechanisms of both sterol-dependent and bexarotene-induced remyelination.

---

Oligodendrocytes (OLs) myelinate axons and provide metabolic support for neurons within the central nervous system (CNS), which facilitates neuronal health and function, as well as the rapid saltatory transmission of action potentials^1^. In addition to these familiar essential roles, OLs have emerged as playing important roles in the context of memory and learning^2–5^. Adaptive myelination, defined by changes in OLs and the myelin they form in response to the activity of associated neurons, serves to reinforce active pathways and contributes to learning^5^. The loss of OLs and myelin results in failed neuronal function and, in the absence of remyelination or myelin repair, axonal degeneration. In response to demyelination, OL progenitor cells (OPCs)^6, 7^, a highly abundant cell population present throughout adulthood in the mammalian central nervous system^8^, become activated to drive the regenerative process of remyelination^9, 10^. Remyelination involves and is defined by the local differentiation OPCs, which leads to the generation of new functional myelinating OLs^11–14^.

Multiple sclerosis (MS) is a debilitating autoimmune disease that is characterized by episodes of focal inflammatory and immune responses that target OLs, which leads to the primary demyelination of axons, an inability to adequately compensate for myelin loss^15^, neuronal dysfunction and ultimately axonal loss^16, 17^. Despite impressive beneficial impacts on disease severity and relapse frequency, immuno-modulatory drugs in isolation ultimately fail in the context of progressive forms of MS disease^18^. In this context, effective OPC differentiation- and remyelination-inducing agents, have significant potential as complementary drugs to immune-targeting therapies for the treatment of progressive forms of MS, as well as diverse other forms of demyelinating disease^11, 13, 19, 20^. Based on the abundant numbers of OPCs that have been observed to be present in some MS lesions^21, 22^, it is believed that inhibited OPC differentiation can play a causative contributing role in disease progression^23–27^. Experimental evidence also clearly indicates that remyelination efficiency deteriorates with age, but that failure in remyelination is not a result of an inherent limitation of an aged stem cell population, but rather as a result of extrinsic factors^28–30^.

Cell-based phenotypic screening has enabled the discovery of OPC differentiation-inducing agents and also the identification of new classes of remyelination-related therapeutic targets^31–33^. Using high-throughput cell-based chemical screening approaches, several research groups have identified and also converged on diverse classes of small molecules that enhance OPC differentiation^31–35^. Based on collective findings surrounding the observed ability of muscarinic 1 receptor (M1R) small molecule antagonists to positively impact OPC differentiation^31, 33^, as well as evidence derived from tissue-specific M1R gene knock-out studies in the context of remyelination^36^, clemastine, a functional M1 receptor antagonist approved as a first-generation anti-histamine, was evaluated in a placebo-controlled Phase II clinical trial to assess its ability to function as a remyelination-inducing agent in relapsing-remitting MS patients^37^. Encouragingly, using a maximally tolerated dose, the study met its primary efficacy endpoint based on impact on full field visual evoked potentials (VEP)^37^, and an increase in myelin water fraction in normal-appearing white matter of the corpus callosum was observed based on 3T MRI data^38^. While this clinical result provides evidence for the ability of the M1R antagonist class to promote functional remyelination in MS patients, as well as a validation of phenotype-based discovery approaches, the magnitude of the effect was modest and dose limiting toxicity likely hinders the ability of clemastine to serve as an optimal agent for the treatment of MS. Based on the relationship of cellular potency to tolerated human steady-state exposure properties^39^, we hypothesize that an alternative M1R antagonist could have a more significant impact on remyelination in patients.

Analysis of transcriptional changes following focal demyelination in the rat CNS, as well as observed expression levels in active human MS lesions, led to the hypothesis that retinoid X receptor gamma (RXRγ) signaling serves as a critical positive regulator of OPC differentiation and remyelination^40^. Consistently, pharmacological inhibition or genetic ablation of RXRγ in cultured cells, as well as RXRγ gene knock-out in mice, was found to negatively impact OPC differentiation and remyelination *in vivo*^40^. A placebo controlled phase 2 clinical study was conducted to assess the safety and efficacy of bexarotene, an RXRγ agonist, as a remyelination-inducing therapy in relapsing-remitting MS patients ^41^. While the study failed to meet its primary efficacy outcome based on an imaging measure of lesion myelin content, statistically significant evidence for a positive impact on remyelination was observed based on reduced full field VEP latency^41^.

Notably, a significant number of small molecules associated with a wide breadth of target classes have been reported to induce OPC differentiation^34, 35^. Evidence for the ability of defined 8,9-unsaturated sterols to facilitate OPC differentiation, which accumulate in response to the inhibition of a defined set of downstream enzymes within the cholesterol biosynthetic pathway (i.e., Cyp51, TM7SF2 or EBP, but not DHCR7 or LSS), provided the basis for an intriguing hypothesis that many OPC differentiation-inducing molecules identified to date, including annotated M1R antagonists, operate by this common mechanism^42, 43^. Importantly, however, the downstream target(s) and mechanism of action for these 8,9-unsaturated sterols in the context of OPC differentiation remains to be elucidated and the universality of this common mechanism remains a topic of debate. Finally, for the developable OPC differentiation-inducing agents that have been identified to date, maximal achievable efficacy level is a significant potential issue that could impact the translation of an optimally efficacious remyelination-inducing therapy. Encouraged by previous findings involving the observed ability of drug combinations involving an endogenous metabolite to synergistically enhance OPC differentiation efficacy levels^44^, and also motivated by unanswered mechanistic questions and controversies within the field, here, we performed pairwise drug combination screens. Our findings provide new potential clinical strategies, as well as mechanistic insights into the role M1R and also LXR in the context of 8,9-unsaturated sterol-and bexarotene-dependent OPC differentiation.

## RESULTS

### Profiling the OL differentiation-inducing activities of M1R antagonists

With the objectives of further examining the relevance and role of M1R in OPC differentiation, as well as potentially identifying an alternative to clemastine that possesses a more favorable predicted therapeutic index, using an imaging-based assay involving induced expression of myelin basic protein (MBP) in primary early postnatal rat optic nerve-derived OPCs^31, 44^, we screened an assembled collection of annotated M1R antagonists and related compounds. Many OPC differentiation-inducing agents were identified that achieved an equivalent or superior level of efficacy to that observed for clemastine (Table 1). Most notably, we identified two approved drugs, doxepin and orphenadrine (Fig. 1a), which were found to maximally induce OPC differentiation at single digit nM concentrations (EC_50_ = 3 nM and 9 nM, respectively, Fig. 1b, c, d), representing an ∼50-fold improvement in potency when compared to clemastine. We selected these agents for further characterization, including mechanism of action studies and pair-wise combinatorial screening. As discussed below, in contrast to clemastine, both of these agents have known tolerated human exposure properties that far exceed that which is predicted to be maximally efficacious in the context of OPC differentiation.

**Table 1.**
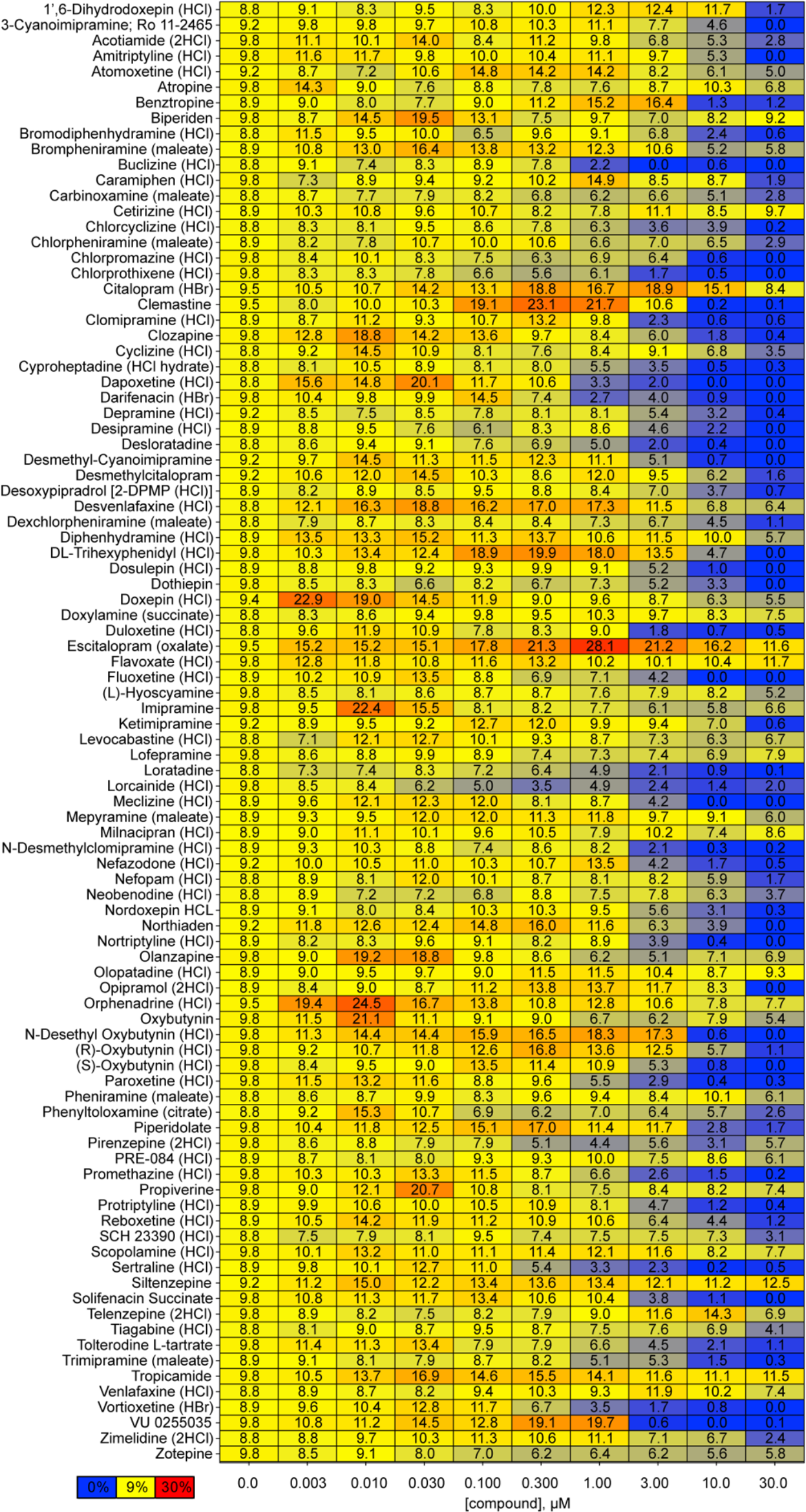
OPC differentiation-inducing activity of muscarinic receptor antagonists. Profiled compounds with reported muscarinic receptor activity. Double gradient heat map showing activity as average (n=3) % of T3-induced MBP+ OLs (blue = 0%; yellow = DMSO baseline; red = largest value).

**Fig. 1.**
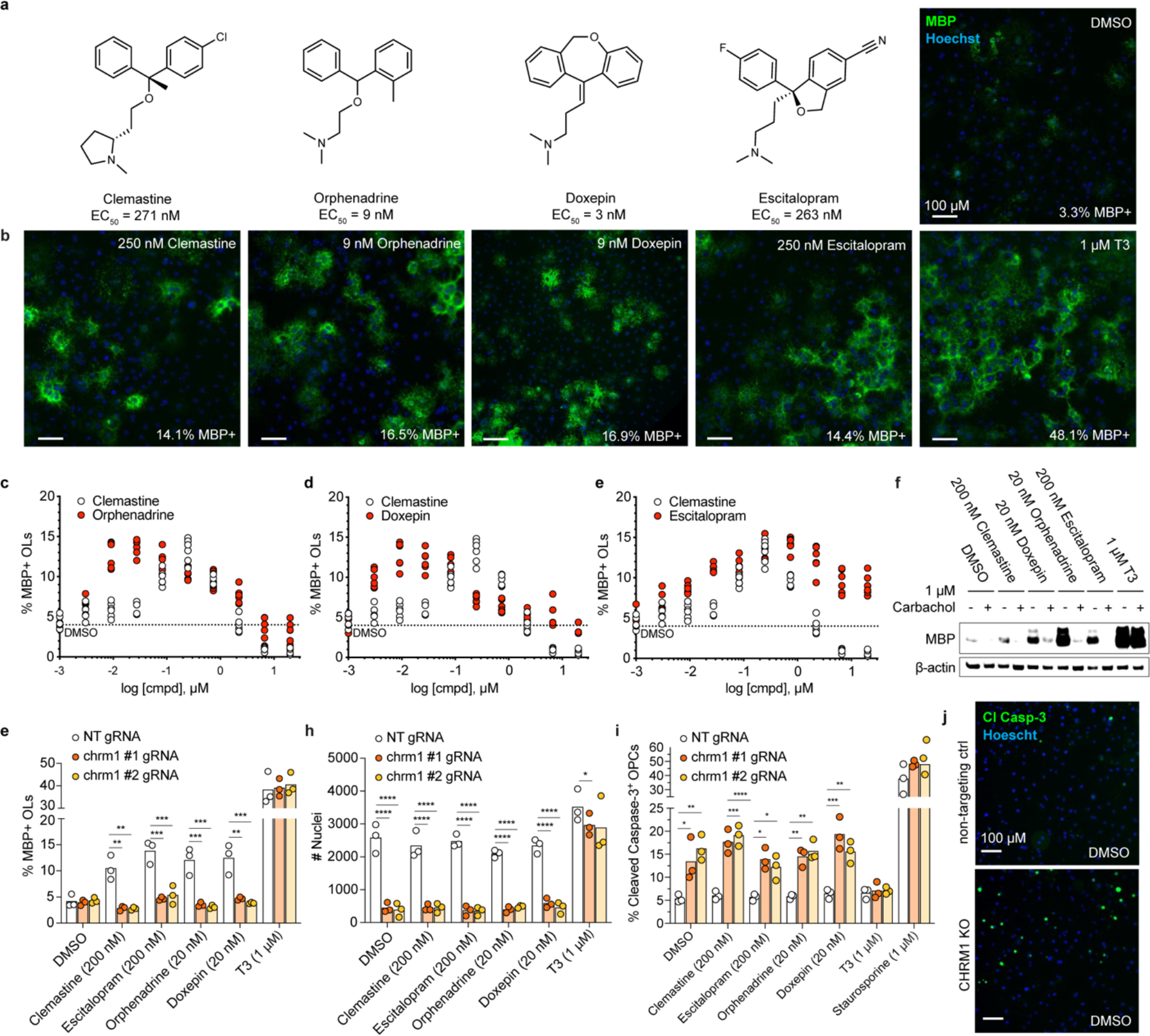
OPC differentiation-inducing activity and mechanisms of M1R antagonists. **a**, Chemical structures and EC50 values for clemastine, orphenadrine, doxepin and escitalopram. **b**, Representative images and % MBP+ values following 6 d of differentiation with clemastine (250 nM), orphenadrine (9 nM), doxepin (9 nM) and escitalopram (250 nM). **c-e**, Dose response curves comparing % MBP+ activity following 6 d of differentiation with clemastine (250 nM) to orphenadrine (9 nM), doxepin (9 nM) or escitalopram (250 nM). **f**, MBP expression following 6 d of differentiation with T3 (1 μM), clemastine (200 nM), escitalopram (200 nM), doxepin (20 nM) or orphenadrine (20 nM) +/- pretreatment with carbachol (1 μM). β-actin was used as internal control. **g**, **h**, %MBP+ cells and total nuclei counts following 6 days of differentiation with clemastine (200 nM), escitalopram (200 nM), orphenadrine (20 nM), doxepin (20 nM) or T3 (1 μM) using OPCs transfected with a non-targeting control gRNA (NT gRNA) or two different M1R-targeting gRNAs (“CHRM1 #1; CHRM1 #2”). **i**, Immunofluorescence-based quantification of cleaved caspase-3-positive OPCs following 72 h of treatment with clemastine (200 nM), escitalopram (200 nM), doxepin (20 nM), orphenadrine (20 nM) or T3 (1 μM). **j**, Representative images showing impact of CHRM1 KO on cleaved caspase-3 expression at 72 h in vehicle-treated OPCs.

### Characterizing the role of M1R in OPC differentiation

Consistent with previous findings^31, 42, 43, 45^, while ∼50% of the compounds we evaluated with characterized M1R antagonistic activity were found to induce some level of OPC differentiation, a remarkable number were observed to be inactive or toxic in this context (Table 1). A notable example is pirenzepine, a selective M1R antagonist^46^, which was observed to lack appreciable OPC differentiation-inducing activity and is cytotoxic at concentrations of ≥100 nM (Table 1). Additionally, on-target potency for M1R does not appear to directly correlate with cell-based potency in OPCs. Reported M1R dissociation constants for clemastine (*K*_i_ = 16 nM)^47^, doxepin (*K*_i_ = 41 nM)^47^, and orphenadrine (*K*_i_ = 38 nM)^47^, are relatively similar and their relative cell-based potency in OPCs differ by >10-fold (Table 1, Fig. 1b-d). To evaluate the mechanistic role of M1R, we examined the impact of carbachol, a muscarinic receptor agonist, on clemastine-, doxepin-, orphenadrine- or escitalopram-induced OPC differentiation. Escitalopram (Fig. 1a) is marketed as a selective serotonin reuptake inhibitor, but is reported to have a modest level of M1R activity (K_i_ = 1.24 uM)^48^ and was found to robustly induce OPC differentiation across a broad concentration range above 100 nM (Fig. 1b, e). As was previously reported for benztropine^31^, carbachol was found to completely ablate the ability of doxepin, orphenadrine or escitalopram to induce OPC differentiation (Fig. 1f, Extended Data Fig. 1a), which supports an essential role for M1R antagonism in the mechanism of action of these compounds and is consistent with tissue-specific M1R gene knock-out studies in the context of remyelination^36^. Importantly, carbachol does not impact the ability of T3 to induce differentiation (Fig. 1f, Extended Data Fig. 1a), indicative of a pharmacological effect that is unique to the M1R-related class of differentiation inducers. In parallel, we examined the impact of Cas9-mediated loss of *chrm1* (M1R encoding gene) on drug-induced differentiation. Electroporation of *chrm1*-targeting gRNAs resulted in complete inhibition of the ability of clemastine, doxepin, orphenadrine or escitalopram to induce OPC differentiation (Fig. 1g, Extended Data Fig. 1b-d), while having no impact on T3-induced differentiation (Fig. 1g, Extended Data Fig. 1c, d). Cas9-mediated *chrm1* loss-of-function in the absence of a differentiation-inducing agent did not result in the induction of OPC differentiation (Fig. 1g), as might be anticipated if M1R signaling serves to negatively regulate this process. However, *Chrm1* deletion was found to have a profound impact on cell viability in the absence of a differentiation-inducing agent, as well as in the context of M1R-related compound-treated conditions, based on total nuclei count (Fig. 1h). In contrast, *Chrm1* deletion had no impact on T3-induced differentiation or cell survival (Fig. 1g, h). Based on these collective observations, we hypothesized that M1R signaling prevents differentiation by maintaining OPCs in a proliferative state and that depending on the tone and strength of pathway inhibition, OPCs are either primed to undergo differentiation or activate programmed cell death. Notably, stem cell fate-related processes are commonly associated with induced apoptosis to ensure accurate lineage specification, and this hypothesis is consistent with observed U-shaped concentration-response curves for this class of agents in this context (Table 1, Fig. 1c-e). To explore this hypothesis, we performed imaging-based quantitative analysis of the apoptosis marker cleaved caspase-3 (cCasp-3). In M1R-deficient cells, increased levels of cCasp-3 were observed under vehicle- or M1R antagonist-treated conditions, but not under T3-treated conditions (Fig. 1i, j, Extended Data Fig. 1e-g). Collectively, these results support a critical role for M1R signaling in the OPC differentiation process and also contributes to an understanding of the mechanism of action of this class of compounds. For this class of agents, M1R antagonism is most likely required but insufficient to induce OPC differentiation.

### Combination-based drug screening for induced OPC differentiation

A recurring observation derived from multiple OPC differentiation assay formats, and across most if not all classes of differentiation-inducing agents, is that the maximal level of achievable efficacy for induced differentiation (i.e., %MBP^+^, MBP^+^ cell count, MBP or marker protein levels, myelin-axon co-localization index, etc.) never reaches the maximal level of the dynamic range that is observed for triiodothyronine (T3) positive control. For example, the maximal level of differentiation efficiency observed for clemastine is typical ∼15-30% of that which is observed for T3, based on either quantitative image analysis of MBP^+^ cells (Table 1) or Western blot analysis of total MBP (Figure 1f). The significant cardiac toxicity liability of thyroid receptor (TR) agonists, as well as associated issues surrounding CNS distribution, currently limits the direct use of existing TR modulating drugs in the context of MS. Inspired by the observed impact of taurine on efficacy levels when paired with diverse classes of OPC differentiation-inducing drugs (i.e., exceeding levels observed for T3 alone)^44^, we sought to determine if defined developable combinations of drugs might result in additive or synergistic effects.

In addition to selecting drugs and tool compounds from reported classes of OPC differentiation-inducing agents, we first completed a screen of the ReFRAME drug repurposing collection (∼12,000 clinical stage or extensively profiled drugs^49^), to ensure alternative and potentially more effective monotherapies had not been overlooked. After validating hits and assessing impact on cell viability, 71 small molecules from the ReFRAME collection were found to have significant OPC differentiation-inducing activity (plate-based z-score ≥3 and ≥15% of T3). Notably, and consistent with reports on the importance of vitamin D receptor (VDR) in the process of OPC differentiation^50, 51^, our screen identified several vitamin D analogues (Extended Data Fig. 2a-d), including calcipotriol, dihydrotachysterol, eldecalcitrol, and tacalcitol. For example, the synthetic vitamin D analogue calcipotriol (Fig. 2a) achieves 40-60% of the efficacy level achieved by T3 across a wide range of concentrations (25 nM to 5 µM; Fig. 2c, Extended Data Fig. 2a). One or more representative tool compounds from each of 11 prioritized hit classes was then selected and subjected to pairwise combination screening to identify additive, synergistic or dose sparing activity (Table 2, Extended Data Fig. 2a, e-l, Extended Data Fig. 3). This resulted in two combinations, each involving a clinically validated class of remyelination-inducing drug (i.e., an M1R antagonist or RXRγ agonist), which were observed to significantly increase differentiation efficacy levels when compared to all monotherapies.

**Fig. 2.**
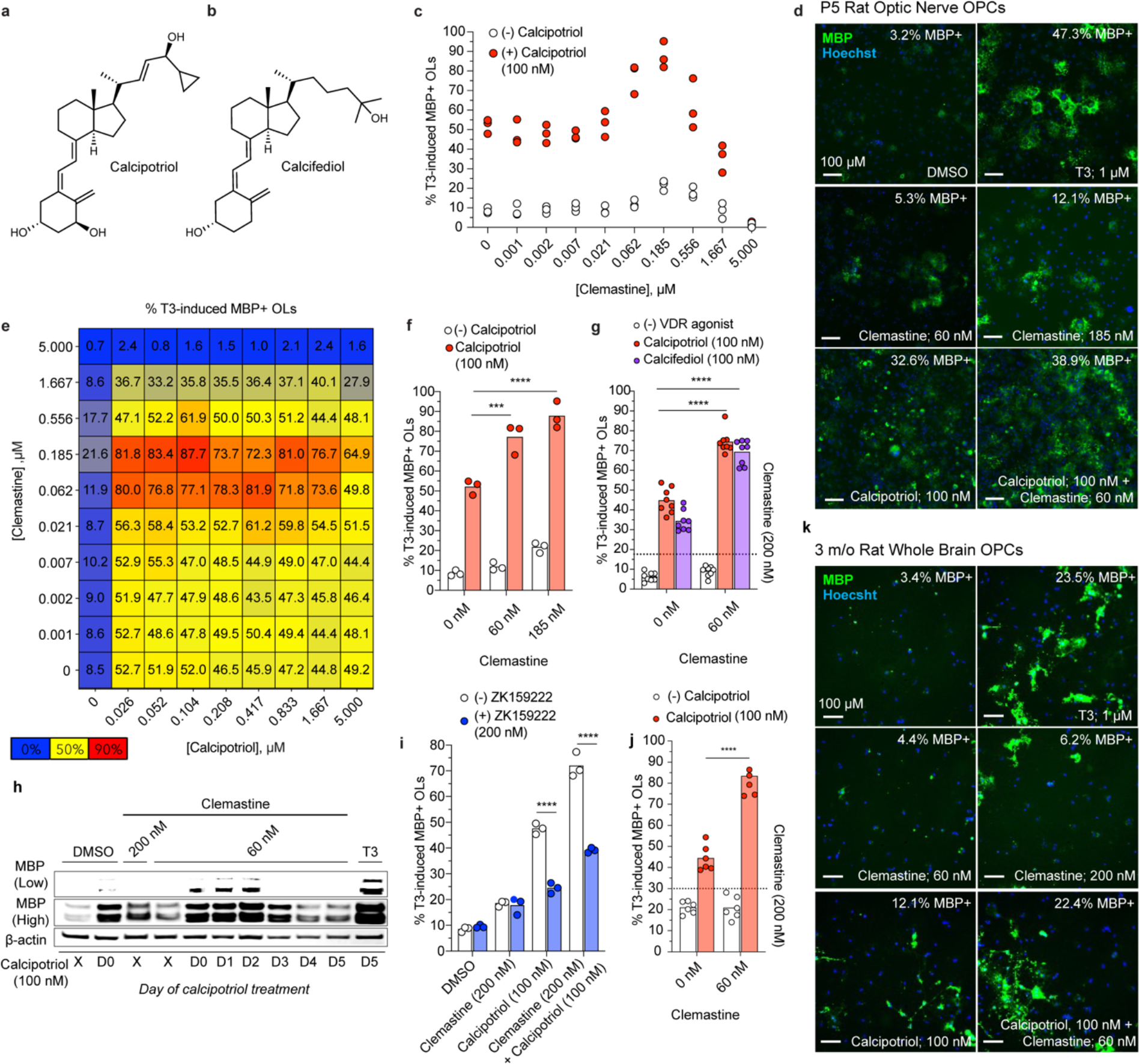
Vitamin D receptor agonism combined with muscarinic receptor antagonism additively enhances OL maturation. **a**, Chemical structures of calcipotriol and (**b**) calcifediol. **c**, OL maturation induced by clemastine (1 nM-5 μM) alone compared to supplementation with bexarotene (500 nM). **d**, Representative images showing MBP expression in P5 rat optic nerve-derived OLs following 6 d treatment with T3 (1 μM), clemastine (60 nM), calcipotriol (100 nM), or calcipotriol + clemastine. **e**, Double gradient heat map showing clemastine +/- calcipotriol activity as average (n=3) % of T3-induced MBP+ OLs (blue = 0%; yellow = optimal % for clemastine; red = largest value). **f**, Comparative impact on % T3-induced MBP+ OLs by clemastine (60 or 185 nM) vs. with calcipotriol (100 nM) added. **g**, Comparative impact on % T3-induced MBP+ OLs by low-dose clemastine (60 nM) vs. with calcipotriol (100 nM) or calcifediol (100 nM) added. **h**, MBP expression at day 6 following the addition clemastine (60 or 200 nM), calcipotriol (100 nM), T3 (1 μM) or a combination of calcipotriol (100 nM, added on days 0-5) and clemastine (60 nM, added on day 0). **i**, Comparative impact on % T3-induced MBP+ OLs by clemastine (100 nM) +/- calcipotriol (100 nM) or ZK 159222 (200 nM). **j**, Image-based quantification of MBP+ OLs following 4 d treatment with calcipotriol (100 nM), clemastine (60 nM) or a combination using OPCs isolated from young rats. Treatment was initiated following 48 h post-isolation recovery. **k,** Representative images showing impact on 3 m/o rat whole brain-derived MBP+ OLs following 4 d treatment with T3 (1 μM), clemastine (60 nM or 200 nM), calcipotriol (100 nM), or calcipotriol + clemastine (60 nM).

**Table. 2.** Profiling of differentiation-inducing compounds for combination studies. Summary of binary combination studies.

| Class | Tool Compound(s) | Partner compound(s) | Observations |
| --- | --- | --- | --- |
| EBP inhibitor | Tasin-1, Tamoxifen | Clemastine, Escitalopram | No effect |
| SERM / ERβ agonist | Tamoxifen, Indazole | Clemastine, Escitalopram | No effect |
| H3R antagonist | Pitolisant, GSK239512 | Clemastine | Additive effect based on total mature cell count |
| HMG-CoA reductase inhibitor | Simvastatin | Clemastine, Escitalopram | No effect |
| M1R antagonist | Clemastine | Escitalopram, Orphenadrine, Doxepin | No effect or potential negative impact |
| PDE inhibitor | Sildenafil | Clemastine, Escitalopram | No effect |
| PPARγ agonist | Pioglitazone | Clemastine | Mildly dose sparing wrt to clemastine |
| RXR agonist | Bexarotene | Clemastine, Pioglitazone, T3, Tasin-1, Tamoxifen | No effect with clemastine; Inhibits T3 or pioglitazone activity; Enhances tasin-1 and tamoxifen activity |
| TR agonist | T3, Sobiterome | Clemastine, Escitalopram | Sparsely additive effect |
| VDR agonist | Calcipotriol | Clemastine, Escitalopram, Orphenadrine, Doxepin | Additive effect with all; Protective with respect to toxicity of M1R antagonist |
| γ-secretase inhibitor | Semagacestat | Clemastine | Negative impact |

### VDR agonists enhance the OPC differentiation activity of M1R antagonists

Binary agent combinatorial screening revealed that addition of the VDR agonist calcipotriol to clemastine has a significant positive impact on OPC differentiation efficacy levels, when compared to the maximal efficacy associated with the corresponding single agent treatment conditions. Based on quantitative immunofluorescent analysis of %MBP^+^ cells, co-treatment with these compounds results in ∼80-90% of the activity that is observed using T3, compared to a maximal level of ∼20% for clemastine or ∼50% for calcipotriol as single agents, respectively (Fig. 2c-e). The combination is also somewhat dose sparing with respect to clemastine. At a suboptimal concentration of clemastine (60 nM), optimal efficacy can be achieved when combined with a low concentration of calcipotriol (Fig. 2f). 25-hydroxycholecalciferol (calcifediol, Fig. 2b) is hydroxylated by CYP27B1 in the kidney to form calcitriol^52^, the active form of vitamin D. Interestingly, calcifediol has a near identical activity profile to that of calcipotriol in the context of OPC differentiation, including its dose-sparing and at least additive activity with respect to clemastine (Fig. 2g). This result suggests that OPCs have the inherent capacity to metabolize vitamin D precursors to the active form. The magnitude of the drug combination effect is even more apparent based on Western blot analysis, which also demonstrated that supplementation of a suboptimal concentration of clemastine with calcipotriol is most effective between days 1-3 (Fig. 2h, Extended Data Fig. 4k). Co-treatment with a VDR antagonist (ZK 159222, Extended Data Fig. 4j) inhibits calcipotriol-induced differentiation and reduces the effect of adding calcipotriol to clemastine (Fig. 2i), indicating that on-target VDR-dependent activity is responsible for the observed drug combination effects. This same effect was observed for all M1R antagonists that were evaluated, including doxepin (Extended Data Fig. 4a, d, e) and orphenadrine (Extended Data Fig. 4b, f, g), as well escitalopram (Extended Data Fig. 4c, h, i), which further suggests that this compound induces differentiation via the same M1R-involving mechanism as clemastine, doxepin and orphenadrine. Finally, to ensure that the observed combination effect was not limited to OPCs derived from early postnatal optic nerve, we evaluated the combination using whole brain-derived OPCs isolated from 3-month-old rats. In this system, addition of calcipotriol to a suboptimal concentration of clemastine results in the achievement of ∼90% of the activity that is observed using T3 (Fig. 2j, k).

### Impact of bexarotene-based combinations on OPC differentiation

Pairwise combinations involving the RXR-γ agonist bexarotene were observed to have diverse and profound effects on OPC differentiation. Despite having a minimal impact on OPC differentiation when evaluated as a single agent in the P5 rat optic nerve-derived OPC assay system (Extended Data Fig. 2k), bexarotene was found to inhibit the OPC differentiation-inducing activities of a PPAR-γ agonist (pioglitazone, Extended Data Fig. 5a-c), as well as that of T3 (Extended Data Fig. 5a, b, d, Fig. 3e), the TR agonist reference control. In stark contrast, bexarotene was found to have a synergistic positive impact on OPC differentiation when combined with tasin-1 (Fig. 3a, c, e) or tamoxifen (Fig. 3b, d, e). The beneficial impact of combining these agents was also clearly demonstrated based on Western blot analysis of induced MBP expression (Fig. 3f, g). Tamoxifen has been previously demonstrated to induce OPC differentiation, as well as remyelination *in vivo*^53^, and was selected as a compound that represented both the selective estrogen receptor modulators (SERM) and also the 8,9-unsaturated sterol-inducer classes. While classified as a SERM, like bazedoxifene^54^, tamoxifen has been convincingly proposed to function in the context of OPC differentiation independently of ER via its ability to inhibit 3β- hydroxysteroid-Δ8, Δ7-isomerase (also known as emopamil binding protein, EBP) at relevant concentrations^42^. Tasin-1 is also an EBP inhibitor and served as a tool compound to help elucidate the ability of accumulated 8,9-unsaturated sterol intermediates, derived via inhibition of defined enzymes within the cholesterol biosynthesis pathway, to drive OPC differentiation^42^. Consistent with the proposed ER-independent mechanism of action for tamoxifen, the activity of indazole chloride (Ind-Cl), an ER-β agonist with reported remyelination-inducing activity ^55^, which was also identified in our screen of the ReFRAME collection (Extended Data Fig. 2e), was not observed to be impacted when combined with bexarotene (Extended Data Fig. 5e, f). Additionally, in a microsome-based EBP activity assay, tasin-1 and tamoxifen were found to inhibit EBP activity (Extended Data Fig. 5g), while indazole, which induces OPC differentiation at concentrations above 100 nM (Extended Data Fig. 2e, f), was not found to inhibit EBP activity (Extended Data Fig. 5g). We therefore hypothesized that the observed bexarotene-based synergistic OPC differentiation inducing effect was unique to compounds that function via a primary mechanism involving the accumulation of 8,9-unsaturated sterols.

**Fig. 3.**
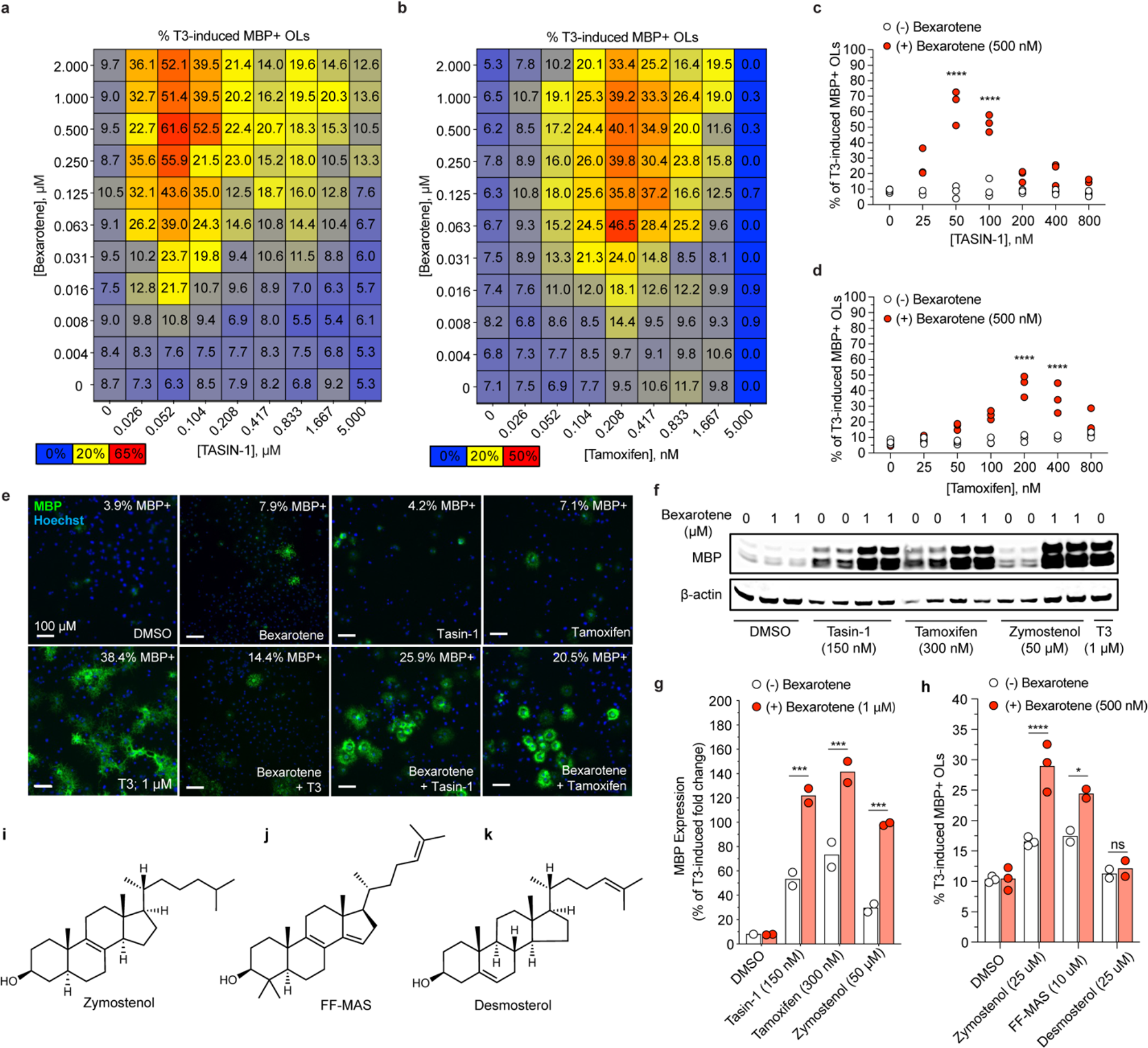
Synergistic differentiation-inducing effect RXR agonism on select compounds that drive accumulation of 8,9-unsaturated sterols. **a**, Double gradient heat map showing bexarotene +/- tasin-1 activity as average (n=3) % of T3-induced MBP+ OLs (blue = 0%; yellow = DMSO baseline; red = largest value). **b**, Double gradient heat map showing Bexarotene +/- Tamoxifen activity as average (n=3) % of T3-induced MBP+ OLs (blue = 0%; yellow = DMSO baseline; red = largest value). **c**, OL maturation induced by tasin-1 (25-800 nM) alone compared to supplementation with bexarotene (500 nM). **d**, OL maturation induced by Tamoxifen (25-800 nM) alone compared to supplementation with bexarotene (500 nM). **e**, Representative images showing impact on MBP+ OLs following 6 d treatment with T3 (1 μM), bexarotene (500 nM), tasin-1 (150 nM), tamixofen (300 nM) or bexarotene (500 nM) + tasin-1 (150 nM) or tamoxifen (300 nM). **f**, MBP expression following 6 d of differentiation with tasin-1 (150 nM), tamoxifen (300 nM), zymostenol (50 μM), or T3 (1 μM). β-actin was used as internal control. **g**, Comparative impact on % T3-induced MBP+ OLs by tasin-1 (150 nM), tamoxifen (300 nM) or zymostenol (50 μM) vs. with bexarotene (1 μM) added. **h**, Comparative impact on % T3-induced MBP+ OLs by zymostenol (50 μM), FF-MAS (10 μM) or desmosterol (25 μM) vs. with bexarotene (500 nM) added. **i-k**, Chemical structures of zymostenol, FF-MAS and Desmosterol.

### Combinations involving 8,9-unsaturated sterols or inducers thereof and bexarotene

A synergistic positive impact on OPC differentiation approaching 50% of T3-induced levels are achieved when >50 nM bexarotene is paired with 50-150 nM tasin-1 (Fig. 3a, c, f, g). These concentrations are consistent with on-target RXRγ agonistic and EBP inhibition activities^42, 47^. Similar levels of efficacy are achieved when >50 nM bexarotene is paired with 100-800 nM tamoxifen, with optimal enhancement at ∼300 nM tamoxifen (Fig. 3b, d, f, g), which is the concentration at which tamoxifen is reported to optimally inhibit EBP^42^. In comparison, as single agent treatments at any concentration, none of these compounds are observed to achieve efficacy levels above ∼12% of that observed for T3 (Fig. 3a, b). Collectively, these results suggested that accumulated 8,9-unsaturated sterols induced by EBP inhibition converge on a mechanism related to RXRγ-dependent OPC differentiation. We therefore assessed the impact of pairing bexarotene directly with defined sterol intermediates from the cholesterol biosynthesis pathway, as well as with small molecule inhibitors of enzymes upstream of EBP (Extended Data Fig. 6a). Consistent with the tasin-1- and tamoxifen-related observations, inhibitors of Cyp51 (i.e., ketoconazole) or Sterol 14-reductase (i.e., amorolfine) were also found to significantly enhance bexarotene-based OPC differentiation (Extended Data Fig. 6b, c). Further, direct addition of zymostenol (Fig. 3i), an 8,9-unsaturated sterol intermediate that accumulates following EBP inhibition (Extended Data Fig. 6a) and was reported to induce OPC differentiation^42^, to bexarotene treatment resulted in a similar synergistic positive impact on OPC differentiation (Fig. 3f, g, h). FF-MAS (Fig. 3j), the related upstream 8,9 unsaturated sterol with demonstrated OPC differentiation-inducing activity^42^, also had a modest impact on bexarotene activity (Fig. 3h). Desmosterol (Fig. 3k), a cholesterol intermediate lacking 8,9 unsaturation and reported to not induce OPC differentiation^42^, was inactive alone and did not impact bexarotene-based OPC differentiation in our assay (Fig. 3h). Notably, results derived using the P5 rat optic nerve-based OPC assay system are completely consistent with the original findings reported by Adams and co-workers related to the ability of 8,9-unsaturated sterols to impact OPC differentiation. Our observed results in the context of bexarotene-based combinations suggested that the ability of RXRγ activation to induce OPC differentiation is dependent on or related to the unknown target responsible for the ability of 8,9 unsaturated sterols to regulate differentiation.

### Liver X receptor modulates accumulated 8,9 unsaturated sterol-based OPC differentiation

RXRs heterodimerize with nuclear receptors (NRs) that have high affinity for their cognate ligands (e.g., TR and VDR) or with lipid activated NRs that have low affinity for their cognate ligands (e.g., PPAR, liver X receptor (LXR), farnesoid-X receptor (FXR) or pregnane-X receptor (PXR)), in either a non-permissive or permissive manner, respectively^56, 57^. Given the observed synergistic effect of combining bexarotene with 8,9 unsaturated sterol-related compounds, the known importance of nuclear receptor signaling in oligodendrocyte lineage specification^24, 40, 50, 58, 59^, and the observed surprising inhibitory impact of bexarotene on NR-based OPC differentiation (i.e., in the context of TR and PPARγ, Extended Data Fig. 5A), we explored known RXR heterodimer partners in the context of 8,9-unsaturated sterol-based OPC differentiation. In contrast to the lack of observed activity of FXR or PXR modulators in this context (Extended Data Fig. 7a, b), LXR modulators were found to significantly impact bexarotene-based effects on OPC differentiation. Addition of an LXR agonist (LXR623, Fig. 4a) resulted in an inhibitory overall impact on differentiation (Fig. 4b), as well as a negative impact on cell viability in some contexts (Fig. 4c). LXR623 was found to completely ablate the synergistic impact of combining bexarotene with an inducer of 8,9-unsaturated sterol accumulation (i.e., tasin-1)(Fig. 4b). Treatment with this LXR agonist had less of an impact on tasin-1-based OPC differentiation in the absence of bexarotene (Fig. 4b, e). Interestingly, the observed negative impact of this LXR agonist on cell viability was also only observed in the context of bexarotene, as this effect was not observed with LXR623 alone or when LXR623 is combined with tasin-1 in the absence of bexarotene (Fig. 4c). Of most mechanistic relevance, a binary combination treatment consisting of an LXR antagonist (GSK2033, Fig. 4d) and bexarotene partially recapitulated the observed beneficial impact of combining bexarotene with a small molecule inducer of 8,9-unsaturated sterol accumulation (i.e., tasin-1, ∼23% versus ∼55% of T3, Fig. 4f, g), and completely recapitulated the effect of combining bexarotene with the relevant corresponding 8,9-unsaturated sterol (i.e., FF MAS, 21.5% versus 24.4% of T3, Fig. 4h). Addition of GSK 2033 did not significantly impact differentiation when added on top of bexarotene already combined with a relevant 8,9-unsaturated sterol (i.e., FF MAS, Fig.4 h), or a small molecule inducer of 8,9-unsaturated sterol accumulation (i.e., tasin-1, Fig. 4f, g). These same relative effects were observed using a structurally orthogonal LXR agonist (i.e., GW3965) or antagonist (i.e., SR9243)(Extended Data Fig. 7e, f). Tasin-1 treatment was not found to impact LXRα/β expression at either the mRNA (Extended Data Fig. 7g), or total protein level (Extended Data Fig. 7h). However, based on confocal image analysis of expression and subcellular localization under various treatment conditions, tasin-1 was observed to increase localization of LXR to the nuclear membrane (Fig. 4i, j, Extended Data Fig.6). Overall, these results suggest that OPC differentiation-inducing 8,9-saturated sterols act as LXR modulators in a manner that is antagonistic and by a potential mechanism involving LXR nuclear trafficking and/or localization.

**Fig. 4.**
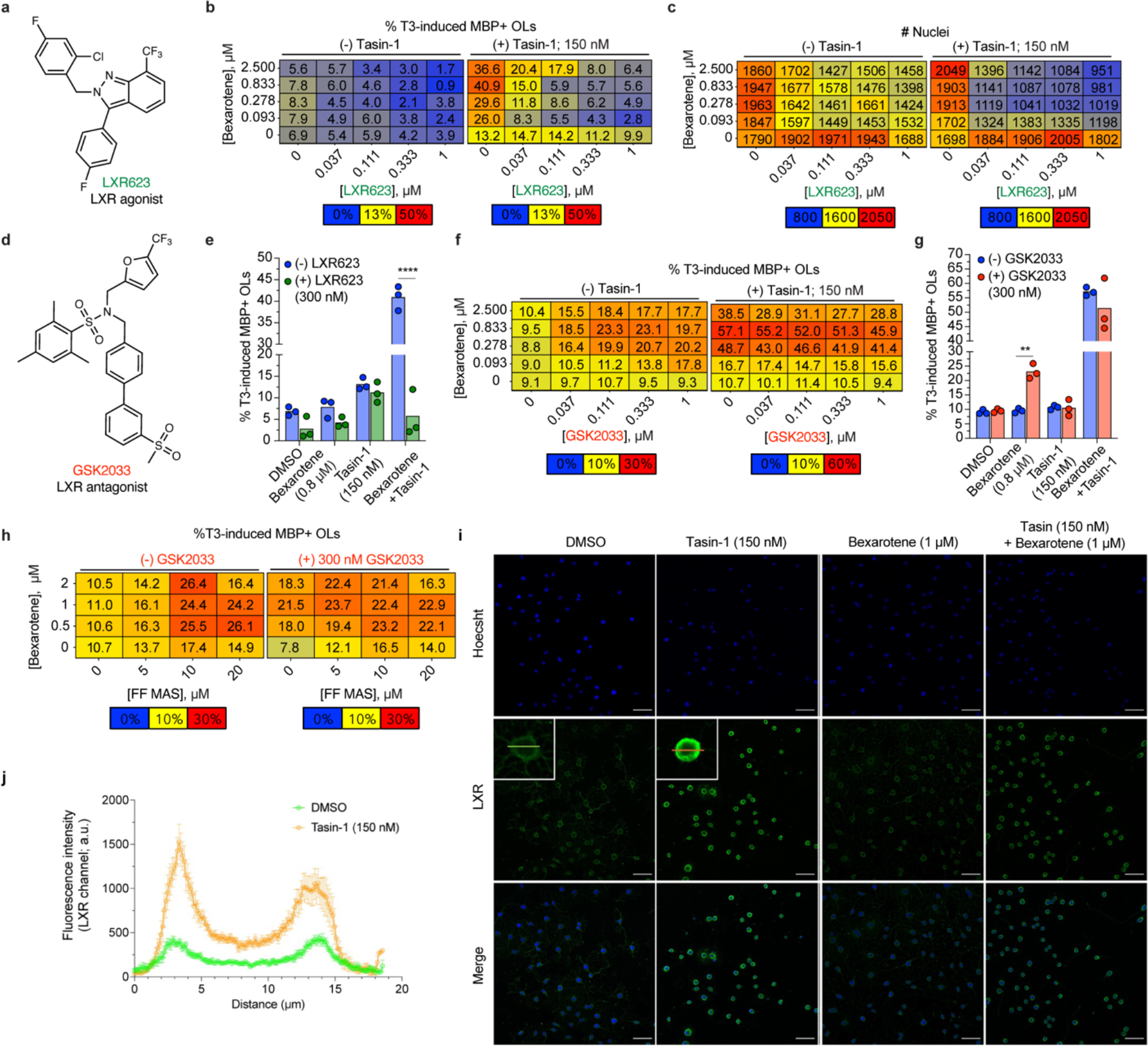
Modulation of select sterol-based mechanisms of remyelination by LXR. **a**, Chemical structure of LXR agonist LXR-623. **b**, Double gradient heat map showing impact of Bexarotene + LXR623 +/- tasin-1 (150 nM) on %T3-induced MBP+ OLs**. c**, Double gradient heat map showing impact of Bexarotene + LXR623 +/- tasin-1 (150 nM) on total nuclei count. **d**, Chemical structure of LXR antagonist GSK2033. **e**, Comparative impact on % T3-induced MBP+ OLs by Bexarotene (800 nM), tasin-1 (150 nM) or a combination of both +/- LXR623 (300 nM). **f**, Double gradient heat map showing impact of Bexarotene + GSK2033 +/- tasin-1 (150 nM) on %T3-induced MBP+ OLs**. g**, Comparative impact on % T3-induced MBP+ OLs by Bexarotene (800 nM), tasin-1 (150 nM) or a combination of both +/- LXR623 (300 nM). **h**, Double gradient heat maps showing impact of GSK 2033 on bexarotene +/- FF-MAS-induced OL maturation, represented as average (n=3) of %T3-induced MBP+ OL. **i**, Representative confocal images (50x) showing LXR expression and localization following 6 d treatment with tasin-1 (150 nM), tamoxifen (300 nM), bexarotene (1 μM), or tasin-1 + bexarotene. Scale bars indicate 50 um. **j**, Line profile showing fluorescent intensities of individual cells (n=10) across an 18.5 um line bisecting each cell (as shown in **i**).

## Discussion

Enabled by the pioneering foundational work of many^1, 6, 7, 9, 11, 13^, breakthrough therapeutic approaches for the treatment of demyelinating diseases involving agents that stimulate the regenerative process of remyelination are on the cusp of becoming a reality. Collective findings derived from genetic analysis and phenotypic screening have provided the basis for clinical trials that have evaluated the ability of repurposed drugs to function as remyelination-inducing agents in the context of MS^41, 60, 61^. The observed evidence for drug-induced remyelination are encouraging and provide proof of concept for the approach^38, 41, 60, 61^. Cell-and patient-based efficacy levels, as well as limitations associated with toxicity concerns, suggest that significant improvements are achievable using alternative agents or approaches. In the ReBUILD trial, Clemastine treatment resulted in a reduction in latency delay of 1.7 ms/eye^37^, as well as an increase in myelin water fraction in the normal appearing white matter of the corpus callosum of clemastine-treated patients^38^. Assuming linear dose-proportional exposure^39^, the predicted steady state concentration of clemastine associated with the maximally tolerated 5.36 mg/kg twice-a-day dose used in the ReBUILD trial is <20 nM, and an optimal concentration of 200 nM is required to achieve <30% of the efficacy observed for T3 in cell-based OPC differentiation assays. In the CCMR One trial^41^, Bexarotene treatment resulted in a reduction in latency delay of 4.75 ms/eye but was associated with tolerability issues at the evaluated dose, and this agent is observed to be only minimally active in our cell-based OPC differentiation assays as a single agent.

In this study we attempted to improve upon these findings and address issues associated with efficacy levels and predicted tolerability, as well as provide potential insights into unanswered mechanistic questions in the field. We first screened a collection of M1R antagonists to find alternatives to clemastine with better predicted human therapeutic indices and performed combination-based drug screening to identify drug pairs that might facilitate improvements in observed efficacy. Two M1R antagonists were identified that induce a maximal level of OPC differentiation at concentrations of <40 nM, compared to 200 nM for clemastine-induced differentiation. Doxepin is a generic widely prescribed medication, which is used to treat conditions ranging from anxiety to sleeping disorder. Doses ranging from 3-300 mg are well tolerated and associated with plasma levels ranging from 3-900 nM^62, 63^. Orphenadrine (norflex) is also a generic medication and is used to treat muscle pain and control motor function in Parkinson’s disease. Doses of 50-100 mg are associated with tolerated human plasma levels that exceed 200 nM^64^. As such, both of these agents have known tolerated human exposure properties that exceed those which are predicted to be maximally efficacious in the context of OPC differentiation, having cell-based potencies of <40 nM and tolerated doses of >100 mg being associated with human steady state exposure levels of >200 nM. In contrast, the steady state human exposure associated with a maximally tolerated dose of 10 mg/day of clemasine is predicted to be ∼20 nM^39^, and a concentration of 200 nM is required to achieve maximal OPC activity in our cell-based system.

Consistent with results from tissue specific M1R knock-out in mouse disease models^65^, pharmacological and genetic results from our study suggest that M1R inhibition serves to facilitate OPC differentiation and is a required component of the mechanism of action for this class of agent in this context. However, the observed lack of activity for many M1R antagonists, including the M1R subtype selective antagonist pirenzepine, suggest that M1R is not the only target responsible for induced OPC differentiation. For this class, M1R inhibition is most likely required but insufficient to drive OPC differentiation, with acetylcholine-based calcium signaling maintaining OPCs in a proliferate state^66, 67^, but more than cell cycle exit being required for efficient differentiation. Unfortunately, we did not identify an agent from our M1R antagonist profiling screen capable of achieving >30% of the efficacy observed using T3. However, pairwise combination screening, using clemastine, doxepin or orphenadrine as an anchor, resulted in the observation that combining any of these agents with a VDR agonist results in efficacy levels that are significantly greater than is observed for members of either class as single agents, and approaching the level achieved using T3. VDR agonists were identified as a robust class of OPC differentiation inducers in our screen of the ReFRAME collection, which is consistent with studies that suggest vitamin D may play a direct role in remyelination by promoting OPC differentiation^50^, and the observed ability of vitamin D (cholecalciferol) supplementation to promote remyelination in a lysolecithin-based rat model^51^. Our results suggest pairing the release of inhibitory signals, using a member of the M1R antagonist class, with the OPC differentiation- and/or maturation-inducing activities of a VDR agonist significantly enhances the overall process of OL formation.

Binary drug combination screening also resulted in the discovery that the ability of bexarotene or tasin-1 to induce OPC differentiation can be synergistically enhanced when these agents are combined. A similar level of benefit was observed when bexarotene was combined tamoxifen, an approved drug. For this latter combination, the cell-based OPC differentiation EC_max_ of bexarotene is ∼100 nM. Assuming dose-proportional exposure, the predicted C_max_ associated with the 300 mg/m^2^ dose of bexarotene used in the CCMR One trial would be 2-10 µM^41, 68^, which approximates the mid-µM levels required to observe cell-based activity in the context of cutaneous T-cell lymphoma (CTCL)^69^, for which bexarotene is licensed and approved. Tamoxifen is a well-established nonsteroidal chemopreventive breast cancer drug, and a 20 mg daily dose of tamoxifen is generally very well tolerated and associated with a steady state plasma concentration of 834 nM^70^, and both of these approved drugs are known to have excellent CNS exposure properties ^70, 71^. Our results suggest that the remyelination inducing activity of bexarotene could be achieved at a lower tolerated dose, and that significant improvements in efficacy might be observed, when combined with tamoxifen or an alternative agent from this mechanistic class. Notably, this defined combination results in an ∼10-fold increase in overall efficacy when compared to the effect of either agent on its own.

Bexarotene was found to have profound and differing effects when evaluated in combination with alternative mechanistic classes of OPC differentiation-inducing agents. Surprisingly, it was found to inhibit TR or PPARγ agonist-induced OPC differentiation. In stark contrast, bexarotene had an additive or synergistic positive impact on the ability of defined 8,9-unsaturated sterol intermediates derived from the cholesterol biosynthesis pathway (i.e., zymostenol, FF-MAS, but not desmosterol), or inhibitors of defined enzymes within the cholesterol biosynthetic pathway that induce their accumulation (i.e., tasin-1, tamoxifen, ketoconazole or amorolfine), to induce OPC differentiation. We used a pharmacology-based approach to explore if this observation might provide insight into how 8,9-unsaturated sterol intermediates of the cholesterol biosynthetic pathway function to induce OPC differentiation, which remains as an outstanding question in the field. Based on the known and observed dominate ability of NR-dependent transcription to regulate the OPC differentiation process, as well as the understood ability of RXRγ to heterodimerize with diverse NR binding partners, we examined if 8,9-unsaturated sterols function by regulating the activity of a known RXRγ binding partner. Uniquely, synthetic LXR antagonists were found to phenocopy the ability of 8,9-unsaturated sterols, or inducers thereof, to combine with bexarotene to enhance induced OPC differentiation. Supportive of a redundant LXR-based mechanism, co-treatment of LXR antagonist with tasin-1 did not further enhance the bexarotene/tasin-1 combination effect. Further implicating LXR as a functional target of 8,9-unsaturated sterols in this context, tasin-1 was found to significantly induce LXR localization to the nuclear membrane of OPCs.

The ability of OPC differentiation-inducing 8,9-unsaturated sterol intermediates to function as LXR modulators would not be unexpected, given that naturally occurring oxysterols, derived downstream of cholesterol from position specific oxidation of the sterol side chain, bind to LXR with high affinity and are thought to function as the canonical endogenous ligands^72, 73^. Importantly, our findings suggest that the OPC differentiation activity of specific 8,9-unsaturated sterols is derived from their ability to inhibit LXR, rather than from their ability to act as agonists. Indeed, synthetic LXR agonists ablated the bexarotene/tasin-1 combination effect. This hypothesis would be consistent with the finding that desmosterol acts as an LXR agonist and lanosterol does not^73^, while the former is inactive and the latter active in the context of OPC differentiation^42^. If 8,9-unsaturated sterols function by inhibiting LXR activity to initiate OPC differentiation, in the context of our collective findings involving bexarotene-based combinations, it is tempting to speculate how diverse NRs orchestrate the overall process with RXRγ serving a rheostat-like function that either enhances or inhibits the initial and secondary stages of OPC differentiation. RXRγ forms heterodimers with NRs that promote differentiation (e.g., TR, VDR, PPARγ) and also with LXR^56, 57^. If LXR-dependent gene expression, or the formation of LXR repressive complexes, serves to inhibit the initiation of differentiation, an RXRγ agonist would enhance these inhibitory signals. This would be consistent with the observed ability of bexarotene to inhibit the differentiation-inducing activities of T3 and pioglitazone. In contrast, the presence of a functional inhibitor of LXR would release this inhibitory signal, resulting in some level of induced differentiation, and when paired with an RXRγ agonist would enable RXRγ to potentiate the pro-differentiation activities of alternative heterodimers (e.g., TR, VDR, PPAR), resulting in additive or synergistic positive effects. This would be consistent with our observation that a significant positive impact on OPC differentiation occurs when bexarotene is combined with agents associated with the 8,9-unsaturated sterol mechanism or an LXR antagonist.

A vast majority (70-80%) of the cholesterol found in the CNS is sequestered within the lipid-rich membranes of myelin^74^, and cholesterol makes up >25% of myelin total lipid content^75^. Further, within the brain cholesterol originates primarily from *de novo* synthesis, as an intact BBB prevents transport from the periphery, and its local metabolism is thought to play critical cell type-specific roles during demyelination and remyelination^76^. As such, it is not surprising that cholesterol sensing mechanisms would play essential roles regulating the processes of OPC differentiation and oligodendrocyte maturation. LXR/RXR functions as a sensor of intracellular cholesterol concentration to mediate efflux. LXRα/μ double knock-out was found to alter myelin structure and myelin gene expression in the cerebellum^77^, and LXR agonists were reported to enhance myelin gene expression and oligodendrocyte maturation^77^. Our results implicate LXR as the target responsible for the OPC differentiation-inducing activity of defined 8,9-unsaturated sterols, but suggest that these intermediates act as a functional antagonists. It is possible that LXR plays multiple roles at different stages of differentiation and maturation, with the initial accumulation of cholesterol intermediates triggering an initiation of OPC differentiation, and subsequent accumulation of downstream oxysterols serving as a signal to initiate gene expression required for maturation of the pre-myelinating cell state and the induction of myelination.

**Extended Data Fig. 1.**
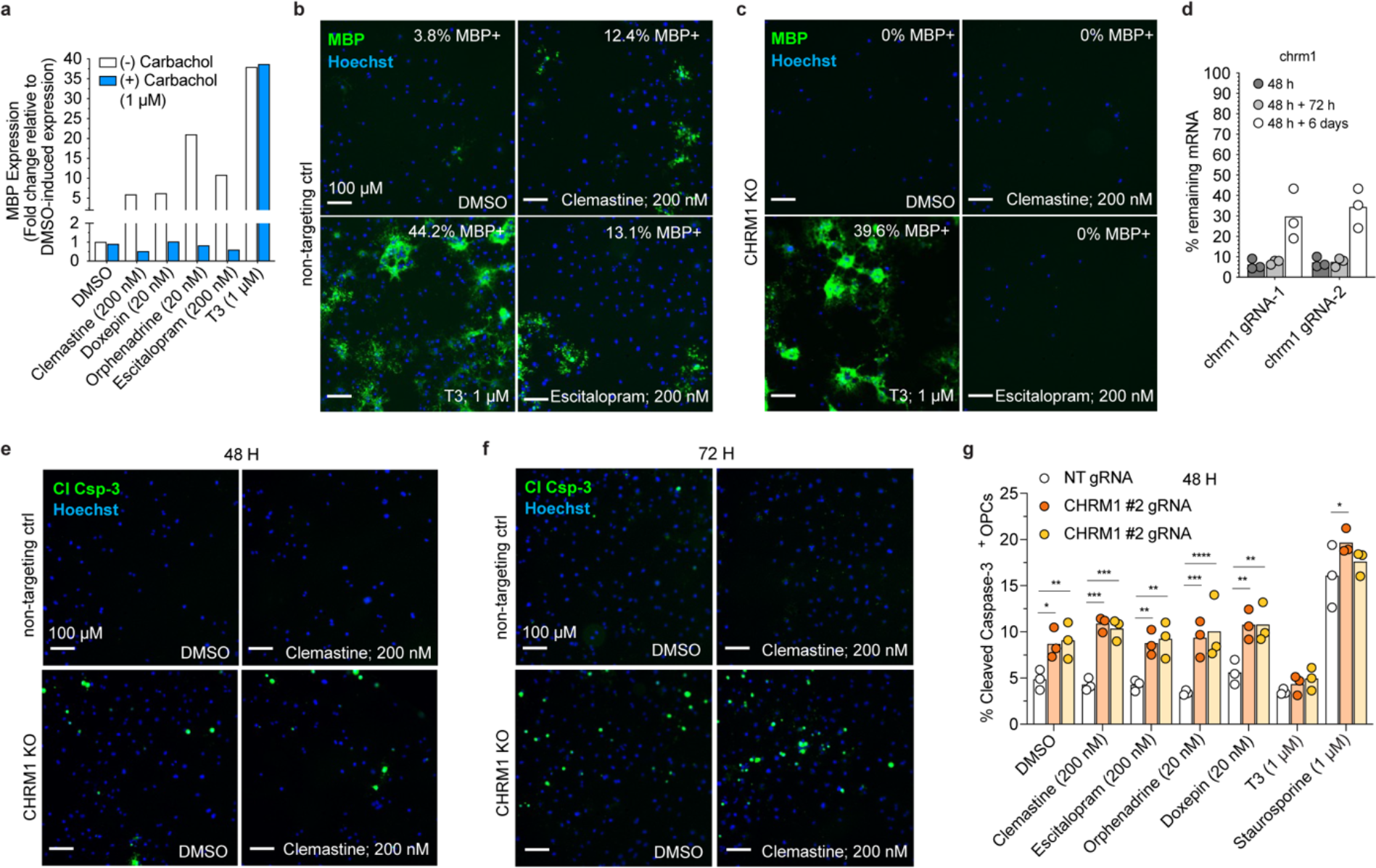
Impact of Cas9-mediated CHRM1 KO on MBP+ or Cleaved Caspase-3+ cells. **a**, Western blot analysis of MBP expression following 6 d of differentiation with T3 (1 μM), clemastine (200 nM), escitalopram (200 nM), doxepin (20 nM) or orphenadrine (20 nM) +/- pretreatment with carbachol (500 nM or 3 μM). Blot shown in Figure 1f. **b**, Representative images of impact of non-targeting control gRNA on clemastine, escitalopram or T3-induced differentiation. **b**, Representative images of impact of CHRM1 KO on clemastine (200 nM), escitalopram (200 nM) or T3 (1 μM)-induced differentiation. **c**, Representative images of impact of CHRM1 KO on clemastine, escitalopram or T3-induced. **d**, Genetic knockout efficiency of two different M1R-targeting gRNAs (“CHRM1 #1; CHRM1 #2”) 48 h after transfection (recovery time), 72 h after recovery time and 6 d after recovery time. **e**, **f**, Immunofluorescence-based quantification of cleaved caspase-3-positive OPCs following 48 h (**e**) or 72 h (**f**) of treatment with clemastine (200 nM). **g**, Immunofluorescence-based quantification of cleaved caspase-3-positive OPCs following 48 h of treatment with clemastine (200 nM), escitalopram (200 nM), orphenadrine (20 nM), doxepin (20 nM) or T3 (1 μM).

**Extended Data Fig. 2.**
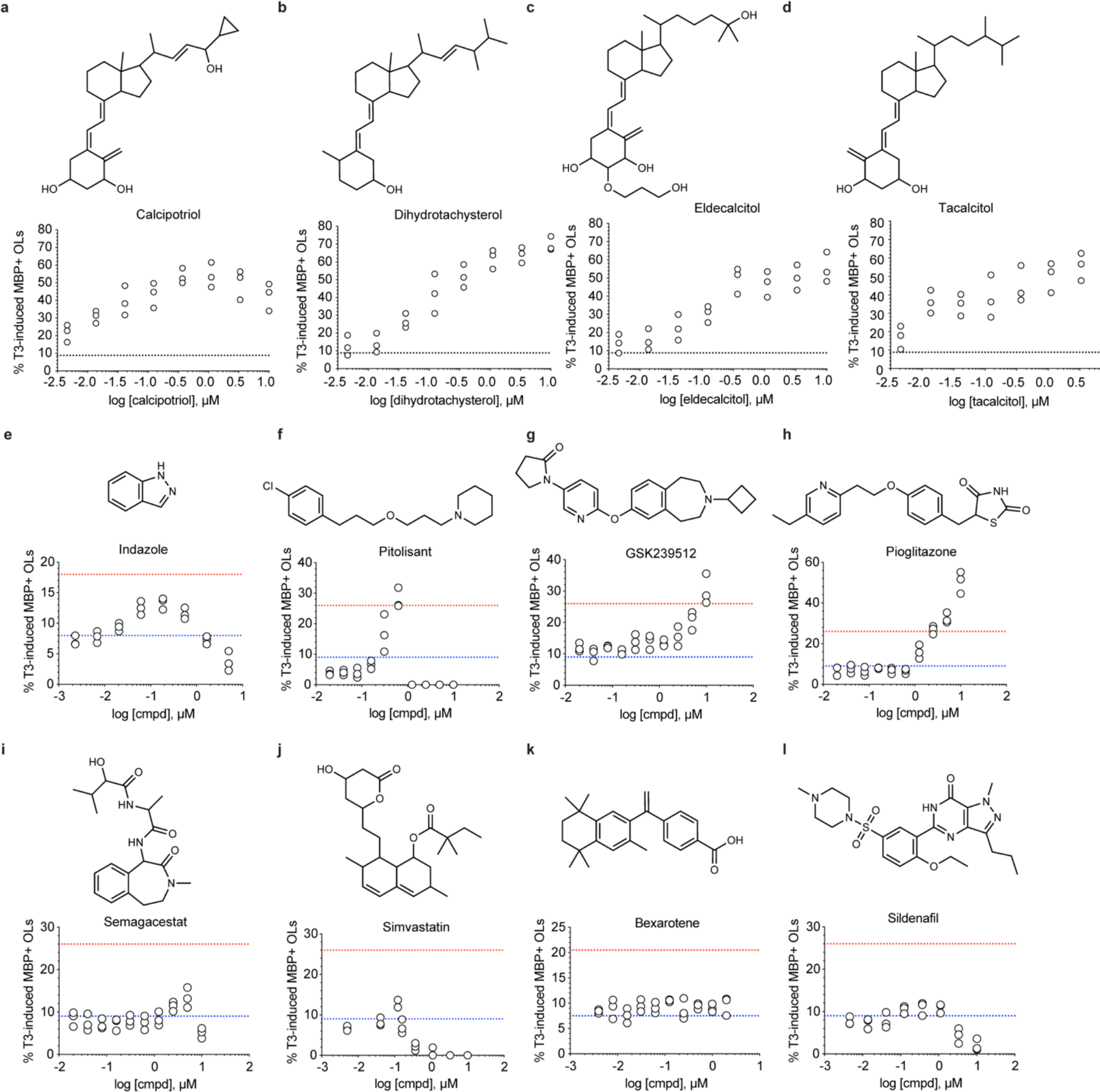
Activity profiles of tool compounds from representative classes. Chemical structure and dose-response activity curves for vitamin D analogues (**a**) calcipotriol, (**b**) dihydrotachysterol, (**c**) eldecalcitol, (**d**) tacalcitol, (**e**) ER-β agonist Indazole, (**f**) H3R agonist Pitolisant, (**g**) H3R agonist GSK 239512, (**h**) PPARγ agonist Pioglitazone, (**i**) γ-secretase antagonist Semagacestat, (**j**) HMG-CoA reductase antagonist Simvastatin, (**k**) RXRγ antagonist Bexarotene and (**l**) PDE inhibitor Sildenafil.

**Extended Data Fig. 3.**
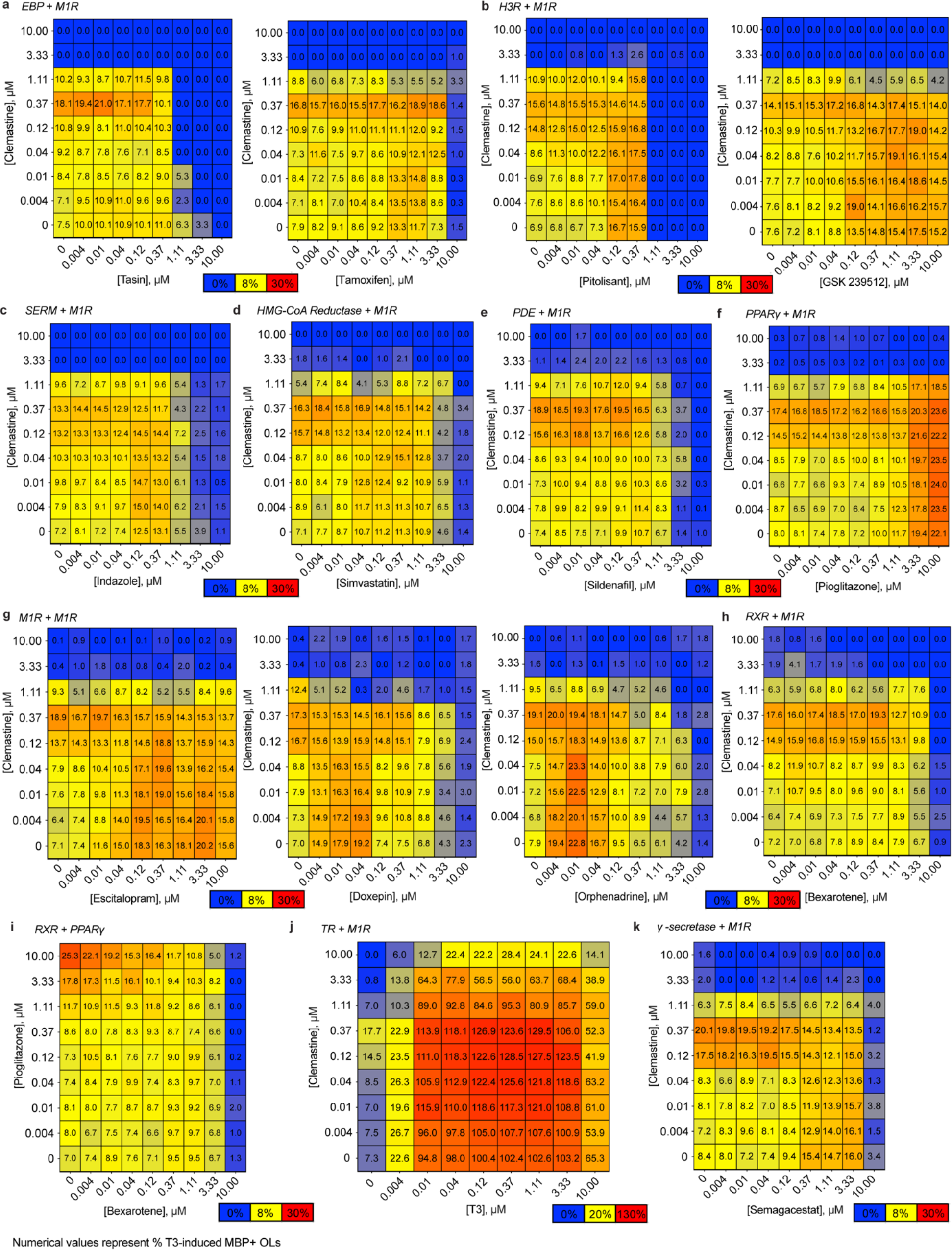
Binary combination studies of tool and partner compounds. **a**, Double gradient heat maps showing M1R antagonist clemastine +/- EBP inhibitors tasin-1 or tamoxifen-induced activity, represented as average (n=3) % of T3-induced MBP+ OLs (blue = 0%; yellow = 8%; red = 30%. **b**, Double gradient heat maps showing clemastine +/- H3R antagonists pitolisant or GSK 239512-induced activity, represented as average (n=3) % of T3-induced MBP+ OLs (blue = 0%; yellow = 8%; red = 30%. **c**, Double gradient heat map showing clemastine +/- ERβ antagonist indazole-induced activity, represented as average (n=3) % of T3-induced MBP+ OLs (blue = 0%; yellow = 8%; red = 30%). **d**, Double gradient heat map showing clemastine +/- HMG-CoA Reductase inhibitor simvastatin-induced activity, represented as average (n=3) % of T3-induced MBP+ OLs (blue = 0%; yellow = 8%; red = 30%). **e**, Double gradient heat map showing clemastine +/- PDE inhibitor inhibitor sildenafil-induced activity, represented as average (n=3) % of T3-induced MBP+ OLs (blue = 0%; yellow = 8%; red = 30%. **f**, Double gradient heat map showing clemastine +/- PPARγ agonist pioglitazone-induced activity, represented as average (n=3) % of T3-induced MBP+ OLs (blue = 0%; yellow = 8%; red = 30%). **g**, Double gradient heat maps showing clemastine +/- partner M1R antagonists escitalopram, doxepin or orphenadrine-induced activity, represented as average (n=3) % of T3-induced MBP+ OLs (blue = 0%; yellow = 8%; red = 30%). **h**, Double gradient heat map showing clemastine +/- RXR agonist bexarotene-induced activity, represented as average (n=3) % of T3-induced MBP+ OLs (blue = 0%; yellow = 8%; red = 30%). **i**, Double gradient heat map showing PPARγ agonist pioglitazone +/- RXR agonist bexarotene-induced activity, represented as average (n=3) % of T3-induced MBP+ OLs (blue = 0%; yellow = 8%; red = 30%). **j**, Double gradient heat map showing clemastine +/- TR agonist T3-induced activity, represented as average (n=3) % of 1 μM T3-induced MBP+ OLs (blue = 0%; yellow = 20%; red = 130%). **k**, Double gradient heat map showing clemastine +/- γ-secretase inhibitor semagacestat-induced activity, represented as average (n=3) % of 1 μM T3-induced MBP+ OLs (blue = 0%; yellow = 8%; red = 30%).

**Extended Data Fig. 4.**
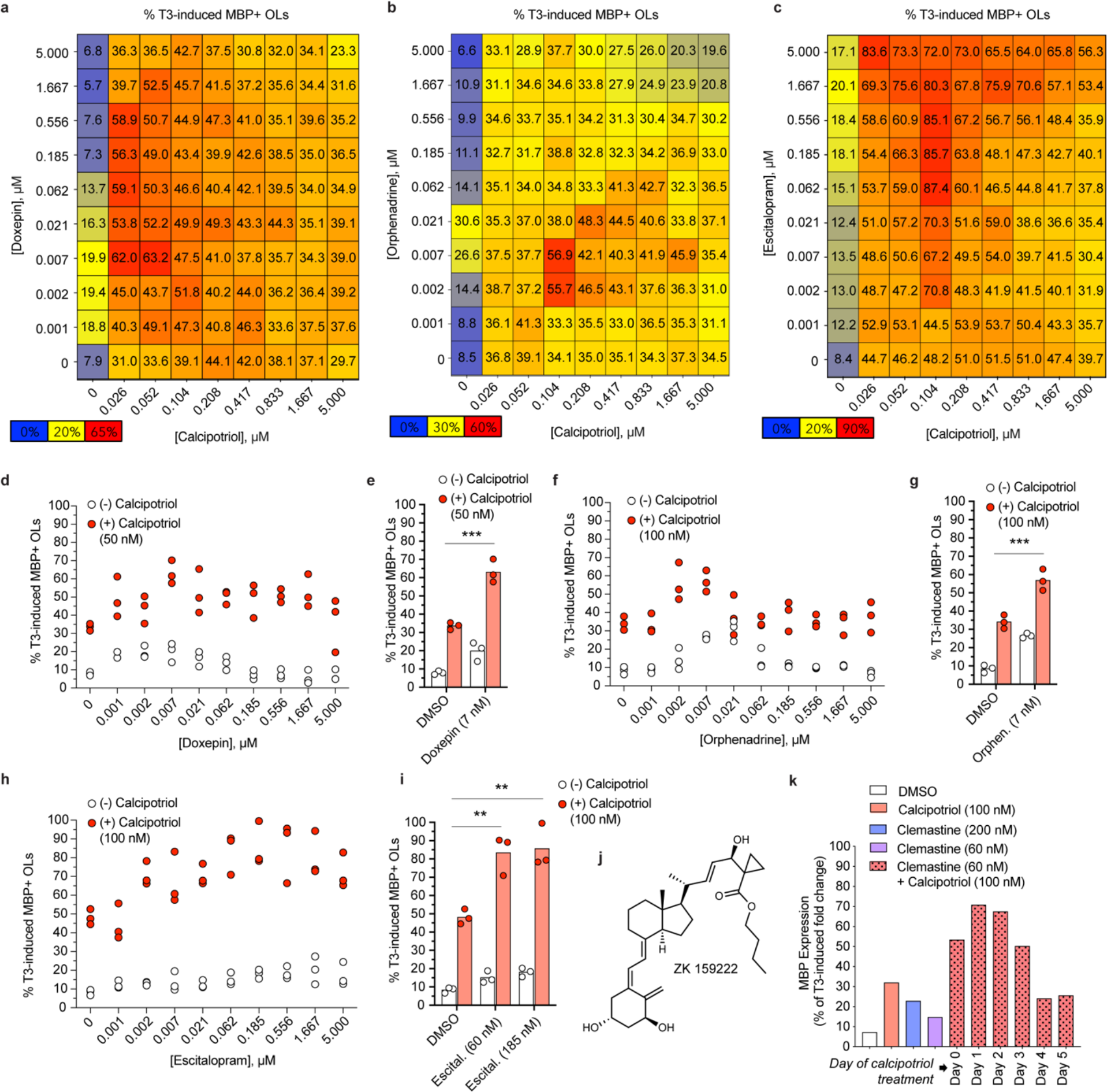
VDR and M1R combinations. **a-c**, Double gradient heat maps showing doxepin (**a**), orphenadrine (**b**) or escitalopram (**c**) +/- calcipotriol activity as average (n=3) % of T3-induced MBP+ OLs (blue = 0%; yellow = optimal % for doxepin; red = largest value). **d**, OL maturation induced by doxepin (1 nM-5 μM) alone compared to supplementation with calcipotriol (50 nM). **e**, Comparative impact on % T3-induced MBP+ OLs by doxepin (7 nM) +/- calcipotriol (50 nM). **f**, OL maturation induced by orphenadrine (1 nM-5 μM) alone compared to supplementation with calcipotriol (100 nM). **g**, Comparative impact on % T3-induced MBP+ OLs by orphenadrine (7 nM) +/- calcipotriol (100 nM). **h**, OL maturation induced by escitalopram (1 nM-5 μM) alone compared to supplementation with calcipotriol (100 nM). **i**, Comparative impact on % T3-induced MBP+ OLs by escitalopram (60 nM or 185 nM) +/- calcipotriol (100 nM). **j**, Chemical structure of VDR antagonist ZK 159222. **k**, Western blot analysis of day 6 MBP expression induced by the addition of calcipotriol (100 nM, added on days 0-5) to clemastine (60 nM, added on day 0). Blot shown in Figure 2h.

**Extended Data Fig. 5.**
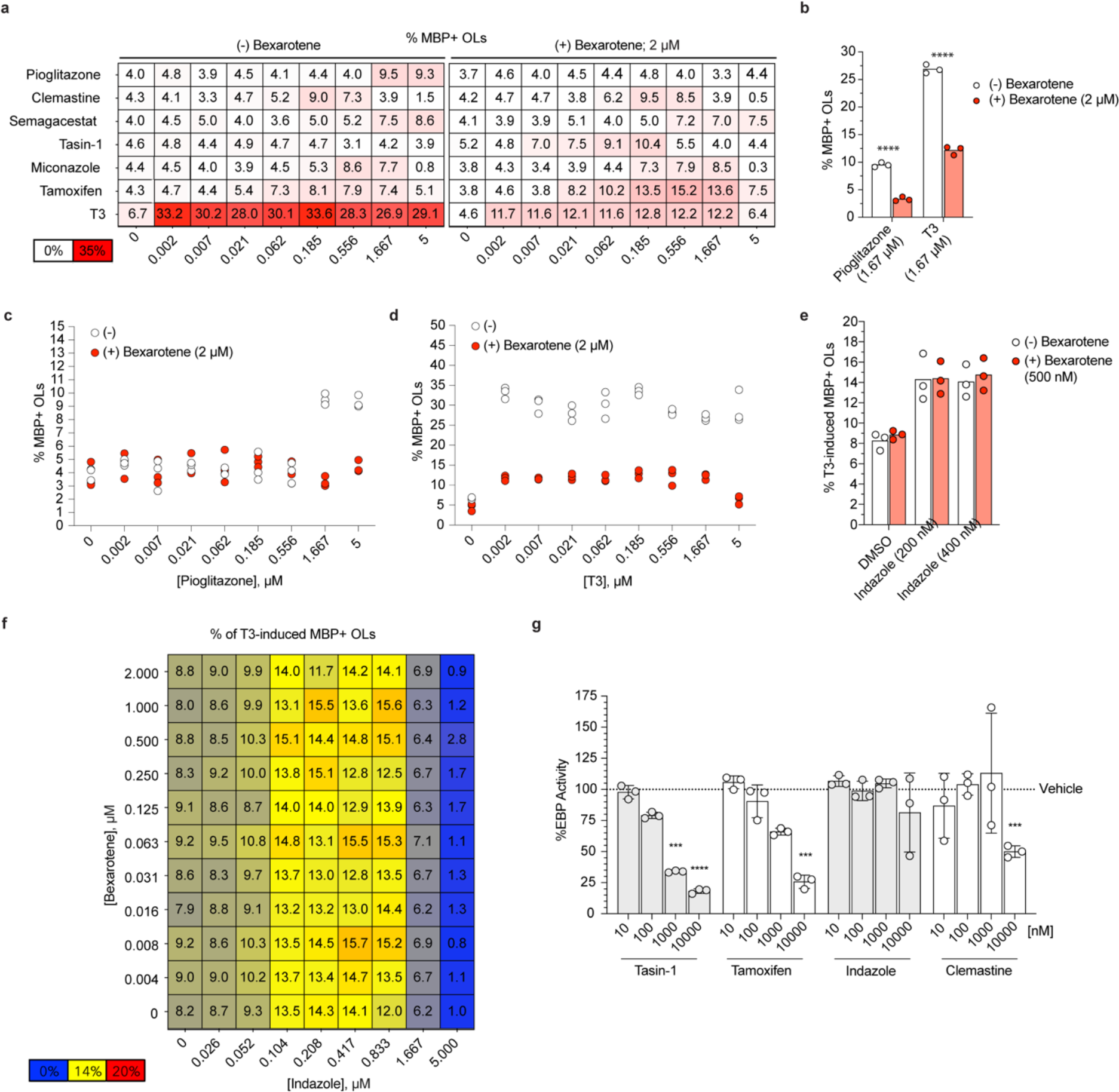
Negative impact of RXR agonist on PPARγ or TR agonists. **a**, Single gradient heat map showing impact of bexarotene (2 μM) addition to pioglitazone, clemastine, semagacestat, tasin-1, miconazole, tamoxifen, or T3 (2 nM to 5 μM). **b**, Comparative impact on % T3-induced MBP+ OLs by pioglitazone (1.67 μM) or T3 (1.67 μM) +/- bexarotene (2 μM). **c**, **d**, OL maturation induced by pioglitazone (2 nM-5 μM; **c**) or T3 (2 nM-5 μM; **d**) alone compared to supplementation with calcipotriol (100 nM). **e**, Comparative impact on % T3-induced MBP+ OLs by indazole (200 nM or 400 nM) +/- bexarotene (500 nM). **f**, Comparative impact on % T3-induced MBP+ OLs by pioglitazone (1.67 μM) or T3 (1.67 μM) vs. with bexarotene (2 μM) added. **g**, %EBP enzymatic activity of tasin-1, tamoxifen, indazole or clemastine (10 nM-10 μM) Significance is based on comparison to avg. normalized values for control (110.4, 110.5, 102.2, 96.5, 92.8, 87.8).

**Extended Data Fig. 6.**
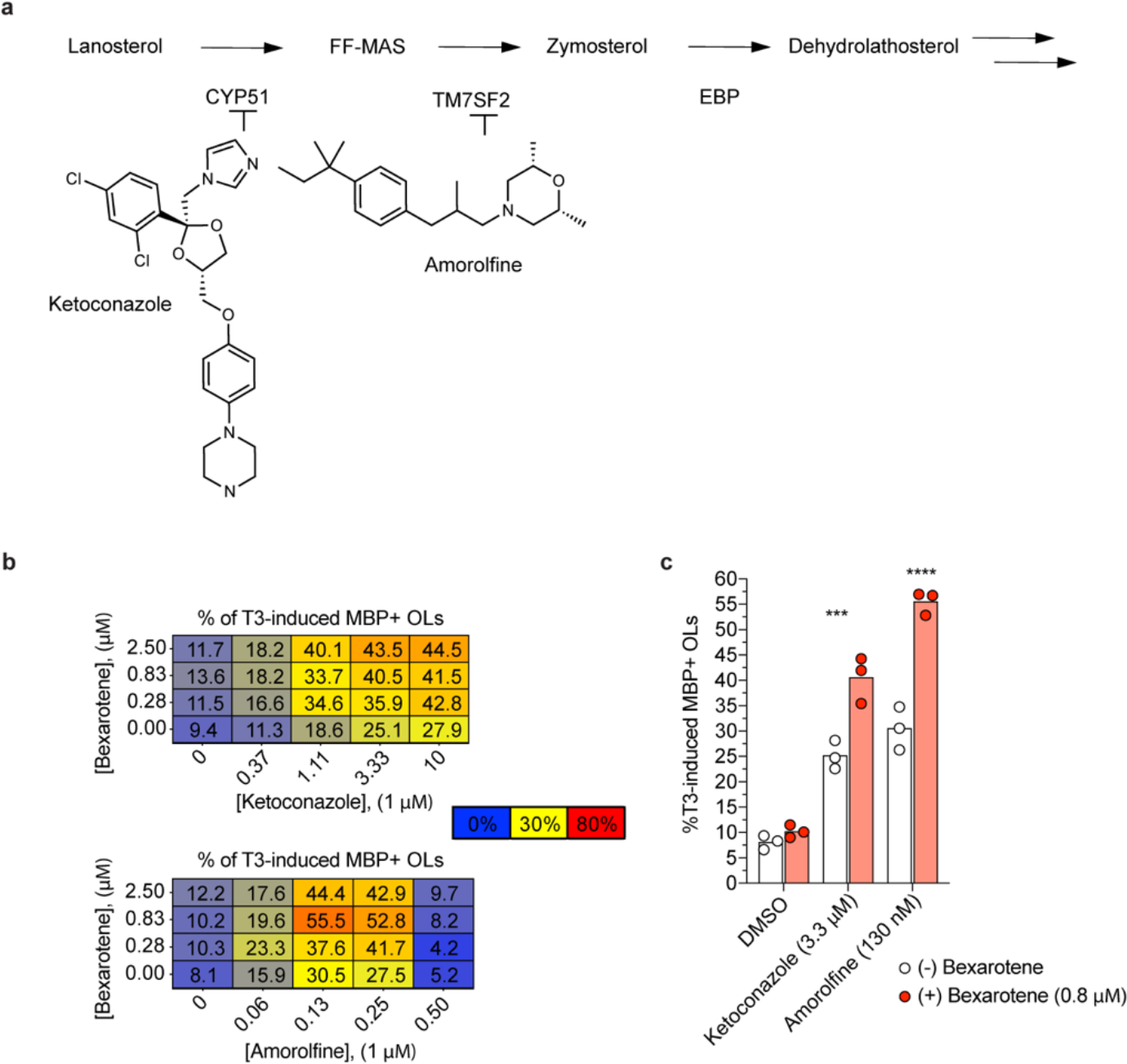
Supplementation of bexarotene to tasin-1, tamoxifen or zymostenol enhances MBP expression. **a**, Abbreviated cholesterol biosynthetic pathway showing inhibition of CYP51 by ketoconazole and TM7SF2 by amorolifine. **b**, Double gradient heat maps showing impact of bexarotene +/- ketoconazole or amorolifine as average (n=3) of %T3-induced MBP+ OL, (blue = 0%; yellow = 30%; red = 80%). **c**, Comparative impact on % T3-induced MBP+ OLs by ketoconazole (3.3 μM), amorolifine (130 nM), +/- bexarotene (0.8 μM).

**Extended Data Fig. 7.**
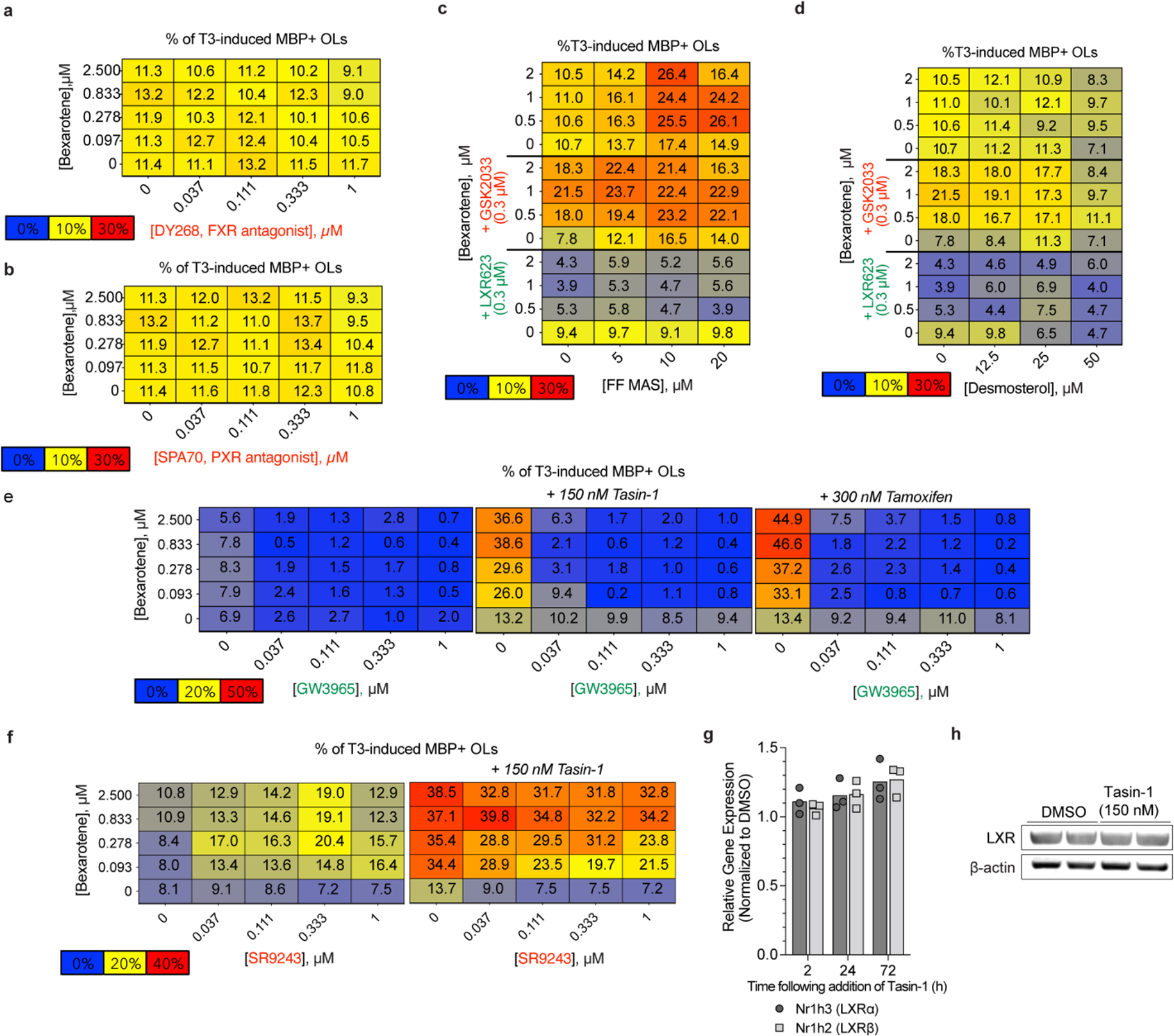
Impact of bexarotene supplementation on other functional inducers of 8,9 unsaturated sterols. **a**, **b**, Double gradient heat maps showing impact of FXR antagonist DY268 (**a**) or PXR antagonist SPA70 (**b**) on OL maturation induced by bexarotene (31 nM to 2.5 μM) + tasin-1 (150 nM), represented as average (n=3) of %T3-induced MBP+ OL, (blue = 0%; yellow = 30%; red = 60%). **c**, Double gradient heat maps showing impact of LXR antagonist GSK 2033 or LXR agonist LXR 623 on bexarotene +/- FF-MAS-induced activity, represented as average (n=3) of %T3-induced MBP+ OL, (blue = 0%; yellow = 10%; red = 30%). **d**, Double gradient heat maps showing impact of LXR antagonist GSK 2033 or LXR agonist LXR 623 on bexarotene +/- Desmosterol-induced activity, represented as average (n=3) of %T3-induced MBP+ OL, (blue = 0%; yellow = 10%; red = 30%). **e**, Double gradient heat maps showing impact of LXR agonist GW3965 on bexarotene +/- tasin-1 (150 nM) or tamoxifen (300 nM)-induced activity, represented as average (n=3) of %T3-induced MBP+ OL, (blue = 0%; yellow = 20%; red = 50%). **f**, Double gradient heat maps showing impact of LXR antagonist SR9243 on bexarotene +/- tasin-1 (150 nM)-induced activity, represented as average (n=3) of %T3-induced MBP+ OL, (blue = 0%; yellow = 20%; red = 40%). **g**, RT-qPCR analysis of relative Nr1h3 (LXRα) or Nr1h2 (LXRβ) expression following 2, 24 or 72 h of treatment with tasin-1 (150 nM). **h**, LXR expression following 6 d treatment with tasin-1 (150 nM).

## Online methods

### Culture and differentiation of P5 rat OPCs

Rat primary optic nerve-derived OPCs were isolated by immunopanning (>99% A2B5^+^) and cultured in poly-D-lysine (10 µg/ml in PBS; Sigma Cat. No. 7280)-coated TC dishes in OPC culture media (Neurobasal Media, Invitrogen) supplemented with B27-without vitamin A (ThermoFisher Cat. No. 12587010), 1x non-essential amino acids, 1x Glutamax, 1x anti-anti, β-mercaptoethanol (all ThermoFisher) and PDGF-AA (50 ng/ml; Peprotech) at 37 °C with 5% CO_2_. The culture medium was replaced every 48h and cells were collected before the confluency reached 60% to maintain a naive state. For differentiation assays, OPCs were seeded at 1K cells per well of a 384-well plate or 200K cells per well of a poly-D-lysine-coated 6-well plate in differentiation media, which is identical to the culture media, but with PDGF-AA supplementation reduced to 2 ng/ml. T3 or 3,3’,5-Triiodo-L-thyronine (Sigma) and DMSO were used as the positive control and negative control, respectively.

### Immunofluorescence-based analysis of MBP expression

For M1R antagonist dose-response evaluation, black, clear-bottom 384-well plates (Greiner) were coated with poly-d-lysine (10 µg/mL in PBS). Clemastine, orphenadrine, doxepin or escitalopram (all purchased from Cayman Chemical) were spotted on plates at 10 doses in differentiation media; three-fold dilutions down from 20 µM. OPCs were then seeded at a density of 1K cells per well, and the plates were incubated at 37 °C with 5% CO2 for 6 d. On day 6, cells were then fixed for 20 min with 4% paraformaldehyde solution (Electron Microscopy Sciences) and stained with anti-MBP antibody (cat. #MAB382, EMD Millipore) in 3% BSA, 0.3% Triton X-100 with overnight incubation at 4 °C. The cells were washed and incubated with secondary antibody (Alexa Fluor 488 goat anti-mouse IgG, Life Technologies) and Hoechst 33342 (Invitrogen) for 1 h at room temperature. The cells were washed and plates were sealed and imaged using a Cellomics Cell Insight imaging reader (Thermo). A 10x lens was used to capture nine images per well at both wavelengths (488 and 365 nm), with each image representing a different unique locations field in a single well. For image analysis, Hoechst-stained nuclei and MBP-positive cell bodies were detected using an algorithm that selects for positive cell bodies and nuclei within a range of fluorescent emission values and sizes as determined by fitting parameters to positive (T3, 1 µM) and negative (DMSO, 0.1%) controls. The number of Hoechst-positive objects were enumerated in experiments where cell counts were required. Numerical results from the analyzed images were analyzed using Prism (Graphpad).

### Western blot analysis

OPCs were plated in basal differentiation medium at 1.7 × 10^5^ cells/well of a 6-well plate (“Day 0”). To evaluate the impact of carbachol on small molecule-induced differentiation, cells were pre-treated with carbachol (2 μM; Sigma) for 6 h. After 6 h incubation with carbachol, clemastine (200 nM), doxepin (20 nM), orphenadrine (20 nM), escitalopram (200 nM) or T3 (1 µM) were added. Plates were incubated at 37 °C with 5% CO2 for 6 d. Following incubation on ice for >20 min and sonication, lysed cells were centrifuged (16,000*g* for 15 min at 4 °C). Total protein was quantified using a BCA analysis, and 25 µg of protein from each sample was denatured by boiling with Bolt LDS Sample Buffer (4x) and Sample Reducing Agent (10x). Proteins were electrophoresed using Bolt 4-12% Bis-Tris gels (Life Technologies) and transferred to a PVDF membrane (cat. # LC2005, Life Technologies). The membrane was blocked with Intercept PBS Blocking Buffer (cat. # 927-76003, LI-COR) and incubated overnight at 4 °C with anti-MBP antibody (cat. #MAB382, Millipore) and anti-β-actin antibody (cat. #sc-47778). Blots were incubated with secondary donkey anti-mouse antibody (IRDye800CW, LI-COR) and imaged using Odyssey CLx and Image Studio (LI-COR). For the calcipotriol supplementation time course, DMSO, clemastine (200 or 60 nM) and T3 (1 µM) were added to differentiation media and calcipotriol was added on each day (0-5). On day 6, cells were washed with cold PBS and collected in ice-cold RIPA buffer containing a protease and phosphatase inhibitor cocktail (Thermo). To evaluate the impact of tasin-1, tamoxifen or zymostenol on total MBP expression by Western Blot, cells were treated with compounds on day 0, and cells collected on day 6 as described above. For

### Genetic deletion of *chrm1*

Invitrogen TrueGuide Synthetic gRNAs (3 guides targeting different regions of each gene) were resuspended in TE buffer at a 100 pmol/ul. A 1:1 molar ratio for Cas9:gRNA was incubated in Buffer R (Neon Transfection Kit, Cat. MPK10096) for 15 minutes at RT. OPCs at 30K cells / µl were added to the complex, mixed and removed using a 10 uL Neon tip. Following electroporation (1600 V, 10 ms, 3 pulses), cells were added to pre-warmed OPC culture media at a density of 150K cells / well of a 6 well plate. Transfected cells were incubated for 48 h. Following a 48 h recovery after transfection (Day 0), the media was replaced with PDGF-depleted differentiation media and the 6-day differentiation assay was initiated.

### Validation of Cas9-mediated genetic knockout by real-time PCR (qPCR)

After 48 h of post-electroporation recovery, OPCs were collected from a single well of a 6-well plate for genetic knockout validation by qPCR. RNA was isolated using an RNeasy Plus Mini Kit (Qiagen) and 1 μg of purified RNA was reverse-transcribed into cDNA (VILO, cat. no. 11755050, ThermoFisher Scientific). Gene expression was assessed using Taqman probes for *chrm1* and *gapdh* (internal control) with the Taqman Universal Mix II (cat. no. 4440038, ThermoFisher) following manufacturer’s instructions. Gene expression was normalized using the delta delta Ct method and was reported as fold change in expression.

### Imaging-based analysis of cleaved caspase-3

In poly-d-lysine-coated black clear bottom 96-well plates (Greiner), cells (transfected with chrm1 gRNAs or a non-targeting control gRNA as described in the previous methods section) were seeded in differentiation media at 5,000 cells per well following their 48 h post-electroporation recovery. Clemastine (200 nM), doxepin (20 nM), orphenadrine (20 nM), escitalopram (200 nM), T3 (1 µM) or staurosporine (1 µM) were added immediately after transferring cells to 96-well plates. The plates were incubated at 37 °C with 5% CO2 for 6 d. On day 6, cells were then fixed for 20 min with 4% paraformaldehyde solution and stained with anti-cleaved caspase-3 antibody (cat. #9961, Cell Signaling Technologies) in 3% BSA, 0.3% Triton X-100 with overnight incubation at 4 °C. The cells were washed and incubated with secondary antibody (Alexa Fluor 488 goat anti-rabbit IgG, Life Technologies) and Hoechst 33342 (Invitrogen) for 1 h at room temperature. The cells were washed and plates were sealed and imaged using the method described above, with a modified algorithm to detect cleaved caspase-3-positive cells.

### Screening of ReFRAME compound library and drug combinations

Using the high-content imaging assay described in previous chapters, ReFRAME compounds were screened at 5 uM, confirmed in triplicate, and confirmed hit compounds (>15% T3-induced activity) were tested in a 12-point dose response format at 10 uM with 3-fold dilutions down. Compounds were transferred from source to destination plates by acoustic droplet ejection (Labcyte Echo). Pairwise screening compounds was performed in matrices of 9-12 doses for each and transferred onto 3 separate 384-well plates. Immunofluorescence staining was quantified based on % MBP^+^ OLs as described in the methods sections of previous chapters. Double-gradient heat maps were generated using Prism to highlight impact of the combinations relative to baseline controls.

### Combination-based screening and evaluation of LXR modulators

Pairwise screening compounds was performed in matrices of 9-12 doses for each and transferred onto 3 separate 384-well plates. Immunofluorescence staining was quantified based on % MBP^+^ OLs as described in the methods sections of previous chapters. Heat maps were generated using Prism to highlight impact of the combinations relative to baseline controls.

### Isolation of rat oligodendrocyte progenitor cells

A 3-month-old rat was euthanized using isoflurane and decapitated as soon as the heartbeat and breathing stopped. The brain was then removed quickly and placed into 50ml conical tube containing 15 to 20ml of ice-cold Hibernate A (cat. HACAMG, Transnetyx). The brain was placed onto 60 mm × 15 mm petri dish containing 2 ml of HBSS without Mg^2+^ and Ca^2+^ (cat. #141750095, GIBCO) and divided along the midsagittal plane. Meninges, the olfactory bulb and white matter were mechanically removed using sterilized forceps. The brain was then cut into approximately 1mm^3^ pieces using a scalpel and rinsed with 15 ml of HBSS without Mg^2+^ and Ca^2+^ and transferred into a 15 ml conical tube. Tissue was collected through centrifuge at 100 g for 1 min at room temperature. Supernatant was discarded and tissue was gently resuspended in 10 ml dissociation solution containing 34 U/ml papain (cat. #LS003126, Worthington) and 40 µg/ml Type IV DNase (cat. #D5025, Sigma-Aldrich) in Hibernate A. The brain tissue was dissociated on the orbital shaker at 50 rpm for 40 min at 35°C. 9 ml Ice cold HBSS without Mg^2+^ and Ca^2+^ was added to the conical tubes to stop digestion and centrifuged at 200 g for 3 min for tissue collection. To obtain a single cell suspension, supernatant was replaced with 5 ml of trituration solution containing 2 mM pyruvate (cat. #11360070, ThermoFisher Scientific) and 2% B27 without vitamin A (cat. #12587010, ThermoFisher Scientific) in Hibernate A. Using a 5 ml serological pipette, tissue was triturated very slowly for 10 times and the supernatant was transferred through a 70 µm cell strainer (cat. #542070, Greiner Bio-One) to a 50 ml conical tube filled with 12 ml of 90% Percoll solution (cat. #17-0891-01, GE Healthcare). The tissue was triturated three more times using fire polished Pasteur pipettes with 2 ml of fresh trituration solution each time. 24 ml of DMEMF12 with HEPES (cat. #11039-021, Gibco) was added to make the final Percoll concentration of 22.5%. Tubes were inverted to mix the solution and centrifuged at 800 g for 20 min at room temperature with no brake. Then the supernatant was aspirated apart from 1∼2 ml of the solution and resuspended in 10ml of HBSS without Mg^2+^ and Ca^2+^ then centrifuged at 300 g for 5 min. After removing the supernatant, pellet was resuspended in 1 ml of red blood cell lysis buffer (cat. #R7757, Sigma-Aldrich) for 90 s. 9 ml of HBSS with Mg^2+^ and Ca^2+^ was added and centrifuged at 300 g for 5 min. The cell pellet was resuspended in 500 µl of wash buffer containing 1x PBS, 2 mM Pyruvate, 0.5% BSA and 2 mM EDTA with 2.5 µg of A2B5 primary antibody (cat. #MAB312, Millipore) for every 10 million cells. The suspension was mixed and incubated at 4°C for 15 min. 7 ml of cold wash buffer was added to the tube and centrifuged at 300 g for 5 min. Pellet was resuspended in 80 µl of wash buffer and 20 µl of rat-anti-mouse-IgM antibody (cat. #130-047-302, Miltenyi) and kept in 4°C for 15 min. 7 ml of cold wash buffer was added to the tube and centrifuged at 300 g for 5 min. The pellet was resuspended in 500 µl of wash buffer and the OPCs were then selected using MS column (cat. #130-042-201, Miltenyi) on MiniMACS Separator (cat. #130-042-102, Miltenyi). MS column was pre-wet with 500 µl of wash buffer before adding cthe ell suspension. After the cell suspension was passed through, the column was washed three times, each with 500 µl of wash buffer. A2B5 selected OPCs are plunged out of the column with 1ml of OPC culture medium composed of 1 mM Pyruvate, 10 µg/ml Insulin (cat. #12585014, ThermoFisher Scientific), and 4% Brightcell SOS Neuronal Supplement (cat. #SCM147, Sigma-Aldrich) in BrainPhys Neuronal Medium (cat. #05790, Stemcell Technologies) supplemented with PDGF-αα (50 ng/ml) and bFGF (20 ng/ml).

### OPC differentiation assay using whole brain-derived rat OPCs

OPCs isolated as described in the previous section were a seeded at 8K cells per well onto poly-d-lysine-coated black clear-bottom 96 well plates. They were given a 48 h recovery period in media containing 1 mM Pyruvate, 10 µg/ml Insulin, and 4% Brightcell SOS Neuronal Supplement in BrainPhys Neuronal Medium supplemented with PDGF-αα (50 ng/ml) and bFGF (20 ng/ml). At 48 h, the recovery media was removed and replaced with OPC differentiation supplemented with B27-without vitamin A, 1x non-essential amino acids, 1x Glutamax, 1x anti-anti, β-mercaptoethanol and PDGF-αα at 2 ng/ml. Cells were treated with clemastine (60 nM, 200 nM), calcipotriol (100 nM), clemastine (60 nM) + calcipotriol (100 nM), or T3 (1 µM). The plates were incubated at 37 °C with 5% CO2 for 4 d. On day 4, cells were then fixed and immunostained for MBP.

### Confocal imaging and analysis of LXR expression

Tasin-1, tamoxifen, bexarotene or bexarotene + tasin-1 or tamoxifen (all purchased from Sigma) were transferred onto black, clear-bottom 384-well plates were coated with poly-d-lysine. OPCs were then seeded at a density of 1,000 cells per well, and the plates were incubated at 37 °C with 5% CO2 for 6 d. On day 6, cells were then fixed for 20 min with 4% formaldehyde solution and stained with anti-LXRalpha/beta antibody (cat. #sc-271064, Santa Cruz Biotechnology) in 3% BSA, 0.3% Triton X-100 with overnight incubation at 4 °C. The cells were washed and incubated with secondary antibody (Alexa Fluor 488 goat anti-mouse IgG, Life Technologies) and Hoechst 33342 (Invitrogen) for 1 h at room temperature. The cells were washed and plates were sealed and imaged using a Cellomics Cell Insight imaging reader (Thermo). Randomly-selected fields from 6 wells selected at random were imaged at 50x magnification using a Zeiss 780 confocal microsope. 10 cells within images from each treatment group were randomly selected for line profile analysis. A line profile analysis of fluorescent intensities was performed by drawing an 18.5 μm line through the center of individual cells (n=10), from approximately 2-4 microns beyond the nuclear membrane on each side using the Fiji image processing platform. Fluorescence intensities over distance (μm) was graphed using Prism 9 software (GraphPad).

### EBP Enzymatic Assay

EBP enzymatic activity was measured with a reported method with modifications.^1^ EBP was obtained through human microsomes (Life Technologies HMMCPL) and diluted to 1 mg/ml assay buffer (100 mM K_2_PO_4_ pH 7.4, 140 mM glucose, 10 mM GSH, 0.5 mM EDTA, 20% Glycerol), then inhibitors were added. After preheating to 37°C, zymostenol (Cayman Chemical) was added at a final concentration of 50 µM, in a final reaction volume of 250 µl, and the reaction was incubated at 37°C for 24h. Reaction was quenched with equivolume 10% KOH in EtOH, internal standard was added (5-α-cholestane, 50 µM, Cayman Chemical), and the aqueous solution was extracted 3×0.5 ml of hexanes to recover sterols. Hexanes were concentrated to dryness under ambient air overnight, reconstituted in 100 µl of CHCl_3_, and analysed using the GC/MS method described below.

### Analysis of sterols by GC/MS

Samples were run on an Agilent 6890 GC coupled to an Agilent 5973 inert MSD operated in scan mode using an FFAP CB 25m × 0.32mm column. Temperature program was 70°C for 5min, ramped at 15°C/min to 250°C, then hold for 20min with a 5 min solvent delay. Helium carrier gas was run at 1.2 ml/min flowrate. Injector port was heated to 250°C, and transfer line to 250°C. Following sample preparation, 1 ul was injected in splitless mode. Ion peaks were integrated to calculate sterol abundance against a metabolite standard curve, and quantitation was normalized to an internal standard. Data was processed using Agilent MassHunter Quantitative Analysis for GCMS, ver B08.00. The following *m/z* ion fragments were used to quantitate each metabolite: Zymostenol (386), lathosterol (386), and 5-α-cholestane (372).

## Notes

### Competing Interest Statement

Frequency Therapeutics, Inc. (Lexington, MA) provided funding as part of a sponsored research agreement to identify M1R antagonist-based drug combinations for the induction of remyelination.

## References

1. Nave, K.A. Myelination and the trophic support of long axons. Nat Rev Neurosci 11, 275–283 (2010).

2. McKenzie, I.A. et al. Motor skill learning requires active central myelination. Science (New York, N.Y.) 346, 318–322 (2014).

3. Gibson, E.M. et al. Neuronal activity promotes oligodendrogenesis and adaptive myelination in the mammalian brain. *Science (New York*, N.Y.) 344, 1252304 (2014).

4. Steadman, P.E. et al. Disruption of Oligodendrogenesis Impairs Memory Consolidation in Adult Mice. Neuron 105, 150–164.e156 (2020).

5. Bechler, M.E., Swire, M. & Ffrench-Constant, C. Intrinsic and adaptive myelination-A sequential mechanism for smart wiring in the brain. Dev Neurobiol 78, 68–79 (2018).

6. Raff, M.C., Miller, R.H. & Noble, M. A glial progenitor cell that develops in vitro into an astrocyte or an oligodendrocyte depending on culture medium. Nature 303, 390–396 (1983).

7. Ffrench-Constant, C. & Raff, M.C. Proliferating bipotential glial progenitor cells in adult rat optic nerve. Nature 319, 499–502 (1986).

8. Dawson, M.R., Polito, A., Levine, J.M. & Reynolds, R. NG2-expressing glial progenitor cells: an abundant and widespread population of cycling cells in the adult rat CNS. Mol Cell Neurosci 24, 476–488 (2003).

9. Smith, K.J., Blakemore, W.F. & McDonald, W.I. Central remyelination restores secure conduction. Nature 280, 395–396 (1979).

10. Prineas, J.W., Barnard, R.O., Kwon, E.E., Sharer, L.R. & Cho, E.S. Multiple sclerosis: remyelination of nascent lesions. Ann Neurol 33, 137–151 (1993).

11. Franklin, R.J. & Ffrench-Constant, C. Remyelination in the CNS: from biology to therapy. Nat Rev Neurosci 9, 839–855 (2008).

12. Tripathi, R.B., Rivers, L.E., Young, K.M., Jamen, F. & Richardson, W.D. NG2 glia generate new oligodendrocytes but few astrocytes in a murine experimental autoimmune encephalomyelitis model of demyelinating disease. J Neurosci 30, 16383–16390 (2010).

13. Franklin, R.J.M. & Ffrench-Constant, C. Regenerating CNS myelin - from mechanisms to experimental medicines. Nat Rev Neurosci 18, 753–769 (2017).

14. Zawadzka, M. et al. CNS-resident glial progenitor/stem cells produce Schwann cells as well as oligodendrocytes during repair of CNS demyelination. Cell stem cell 6, 578–590 (2010).

15. Loma, I. & Heyman, R. Multiple sclerosis: pathogenesis and treatment. Current neuropharmacology 9, 409–416 (2011).

16. Pouly, S. & Antel, J.P. Multiple sclerosis and central nervous system demyelination. Journal of autoimmunity 13, 297–306 (1999).

17. Reich, D.S., Lucchinetti, C.F. & Calabresi, P.A. Multiple Sclerosis. The New England journal of medicine 378, 169–180 (2018).

18. Gholamzad, M. et al. A comprehensive review on the treatment approaches of multiple sclerosis: currently and in the future. Inflamm Res 68, 25–38 (2019).

19. Franklin, R.J. & Gallo, V. The translational biology of remyelination: past, present, and future. Glia 62, 1905–1915 (2014).

20. Feinstein, A., Freeman, J. & Lo, A.C. Treatment of progressive multiple sclerosis: what works, what does not, and what is needed. Lancet Neurol 14, 194–207 (2015).

21. Wolswijk, G. Chronic stage multiple sclerosis lesions contain a relatively quiescent population of oligodendrocyte precursor cells. J Neurosci 18, 601–609 (1998).

22. Chang, A., Tourtellotte, W.W., Rudick, R. & Trapp, B.D. Premyelinating oligodendrocytes in chronic lesions of multiple sclerosis. The New England journal of medicine 346, 165–173 (2002).

23. Hart, I.K., Richardson, W.D., Bolsover, S.R. & Raff, M.C. PDGF and intracellular signaling in the timing of oligodendrocyte differentiation. The Journal of cell biology 109, 3411–3417 (1989).

24. Billon, N., Tokumoto, Y., Forrest, D. & Raff, M. Role of thyroid hormone receptors in timing oligodendrocyte differentiation. Dev Biol 235, 110–120 (2001).

25. Kuhlmann, T. et al. Differentiation block of oligodendroglial progenitor cells as a cause for remyelination failure in chronic multiple sclerosis. Brain : a journal of neurology 131, 1749–1758 (2008).

26. Tokumoto, Y.M., Tang, D.G. & Raff, M.C. Two molecularly distinct intracellular pathways to oligodendrocyte differentiation: role of a p53 family protein. EMBO J 20, 5261–5268 (2001).

27. Fernandez, M. et al. Thyroid hormone administration enhances remyelination in chronic demyelinating inflammatory disease. Proceedings of the National Academy of Sciences of the United States of America 101, 16363–16368 (2004).

28. Ruckh, J.M. et al. Rejuvenation of regeneration in the aging central nervous system. Cell Stem Cell 10, 96–103 (2012).

29. Neumann, B. et al. Metformin Restores CNS Remyelination Capacity by Rejuvenating Aged Stem Cells. Cell Stem Cell 25, 473–485.e478 (2019).

30. Segel, M. et al. Niche stiffness underlies the ageing of central nervous system progenitor cells. Nature 573, 130–134 (2019).

31. Deshmukh, V.A. et al. A regenerative approach to the treatment of multiple sclerosis. Nature 502, 327–332 (2013).

32. Najm, F.J. et al. Drug-based modulation of endogenous stem cells promotes functional remyelination in vivo. Nature 522, 216–220 (2015).

33. Mei, F. et al. Micropillar arrays as a high-throughput screening platform for therapeutics in multiple sclerosis. Nature medicine 20, 954–960 (2014).

34. Caprariello, A.V. & Adams, D.J. The landscape of targets and lead molecules for remyelination. Nat Chem Biol 18, 925–933 (2022).

35. Beyer, B.A. & Lairson, L.L. Promoting remyelination: A case study in regenerative medicine. Curr Opin Chem Biol 70, 102201 (2022).

36. Mei, F. et al. Accelerated remyelination during inflammatory demyelination prevents axonal loss and improves functional recovery. Elife 5 (2016).

37. Green, A.J. et al. Clemastine fumarate as a remyelinating therapy for multiple sclerosis (ReBUILD): a randomised, controlled, double-blind, crossover trial. Lancet 390, 2481–2489 (2017).

38. Caverzasi, E. et al. MWF of the corpus callosum is a robust measure of remyelination: Results from the ReBUILD trial. Proceedings of the National Academy of Sciences of the United States of America 120, e2217635120 (2023).

39. Schran, H.F., Petryk, L., Chang, C.T., O’Connor, R. & Gelbert, M.B. The pharmacokinetics and bioavailability of clemastine and phenylpropanolamine in single-component and combination formulations. J Clin Pharmacol 36, 911–922 (1996).

40. Huang, J.K. et al. Retinoid X receptor gamma signaling accelerates CNS remyelination. Nature neuroscience 14, 45–53 (2011).

41. Brown, J.W.L. et al. Safety and efficacy of bexarotene in patients with relapsing-remitting multiple sclerosis (CCMR One): a randomised, double-blind, placebo-controlled, parallel-group, phase 2a study. Lancet Neurol 20, 709–720 (2021).

42. Hubler, Z. et al. Accumulation of 8,9-unsaturated sterols drives oligodendrocyte formation and remyelination. Nature 560, 372–376 (2018).

43. Allimuthu, D. et al. Diverse Chemical Scaffolds Enhance Oligodendrocyte Formation by Inhibiting CYP51, TM7SF2, or EBP. Cell Chem Biol 26, 593–599.e594 (2019).

44. Beyer, B.A. et al. Metabolomics-based discovery of a metabolite that enhances oligodendrocyte maturation. Nature chemical biology 14, 22–28 (2018).

45. Sax, J.L. et al. Enhancers of Human and Rodent Oligodendrocyte Formation Predominantly Induce Cholesterol Precursor Accumulation. ACS chemical biology 17, 2188–2200 (2022).

46. Carmine, A.A. & Brogden, R.N. Pirenzepine. A review of its pharmacodynamic and pharmacokinetic properties and therapeutic efficacy in peptic ulcer disease and other allied diseases. Drugs 30, 85–126 (1985).

47. Ursu, O. et al. DrugCentral: online drug compendium. Nucleic Acids Res 45, D932–d939 (2017).

48. Sanchez, C., Reines, E.H. & Montgomery, S.A. A comparative review of escitalopram, paroxetine, and sertraline: Are they all alike? Int Clin Psychopharmacol 29, 185–196 (2014).

49. Janes, J. et al. The ReFRAME library as a comprehensive drug repurposing library and its application to the treatment of cryptosporidiosis. Proceedings of the National Academy of Sciences of the United States of America 115, 10750–10755 (2018).

50. de la Fuente, A.G. et al. Vitamin D receptor-retinoid X receptor heterodimer signaling regulates oligodendrocyte progenitor cell differentiation. J Cell Biol 211, 975–985 (2015).

51. Gomez-Pinedo, U. et al. Vitamin D increases remyelination by promoting oligodendrocyte lineage differentiation. Brain Behav 10, e01498 (2020).

52. Bikle, D.D. Vitamin D metabolism, mechanism of action, and clinical applications. Chem Biol 21, 319–329 (2014).

53. Gonzalez, G.A. et al. Tamoxifen accelerates the repair of demyelinated lesions in the central nervous system. Scientific reports 6, 31599 (2016).

54. Rankin, K.A. et al. Selective Estrogen Receptor Modulators Enhance CNS Remyelination Independent of Estrogen Receptors. J Neurosci 39, 2184–2194 (2019).

55. Moore, S.M. et al. Multiple functional therapeutic effects of the estrogen receptor β agonist indazole-Cl in a mouse model of multiple sclerosis. Proceedings of the National Academy of Sciences of the United States of America 111, 18061–18066 (2014).

56. Lefebvre, P., Benomar, Y. & Staels, B. Retinoid X receptors: common heterodimerization partners with distinct functions. Trends Endocrinol Metab 21, 676–683 (2010).

57. Rőszer, T., Menéndez-Gutiérrez, M.P., Cedenilla, M. & Ricote, M. Retinoid X receptors in macrophage biology. Trends Endocrinol Metab 24, 460–468 (2013).

58. Melchor, G.S., Khan, T., Reger, J.F. & Huang, J.K. Remyelination Pharmacotherapy Investigations Highlight Diverse Mechanisms Underlying Multiple Sclerosis Progression. ACS Pharmacol Transl Sci 2, 372–386 (2019).

59. Bernardo, A., Bianchi, D., Magnaghi, V. & Minghetti, L. Peroxisome proliferator-activated receptor-gamma agonists promote differentiation and antioxidant defenses of oligodendrocyte progenitor cells. J Neuropathol Exp Neurol 68, 797–808 (2009).

60. Green, A.J. et al. Clemastine fumarate as a remyelinating therapy for multiple sclerosis (ReBUILD): a randomised, controlled, double-blind, crossover trial. Lancet 390, 2481–2489 (2017).

61. Schwartzbach, C.J. et al. Lesion remyelinating activity of GSK239512 versus placebo in patients with relapsing-remitting multiple sclerosis: a randomised, single-blind, phase II study. J Neurol 264, 304–315 (2017).

62. Weber, J., Siddiqui, M.A., Wagstaff, A.J. & McCormack, P.L. Low-dose doxepin: in the treatment of insomnia. CNS Drugs 24, 713–720 (2010).

63. Leucht, S. et al. Doxepin plasma concentrations: is there really a therapeutic range? J Clin Psychopharmacol 21, 432–439 (2001).

64. Labout, J.J., Thijssen, C., Keijser, G.G. & Hespe, W. Difference between single and multiple dose pharmacokinetics of orphenadrine hydrochloride in man. Eur J Clin Pharmacol 21, 343–350 (1982).

65. Mei, F. et al. Accelerated remyelination during inflammatory demyelination prevents axonal loss and improves functional recovery. eLife 5, e18246 (2016).

66. De Angelis, F., Bernardo, A., Magnaghi, V., Minghetti, L. & Tata, A.M. Muscarinic receptor subtypes as potential targets to modulate oligodendrocyte progenitor survival, proliferation, and differentiation. Dev Neurobiol 72, 713–728 (2012).

67. Weider, M. et al. Nfat/calcineurin signaling promotes oligodendrocyte differentiation and myelination by transcription factor network tuning. Nat Commun 9, 899 (2018).

68. Rizvi, N.A. et al. A Phase I study of LGD1069 in adults with advanced cancer. Clinical cancer research : an official journal of the American Association for Cancer Research 5, 1658–1664 (1999).

69. Zhang, C., Hazarika, P., Ni, X., Weidner, D.A. & Duvic, M. Induction of apoptosis by bexarotene in cutaneous T-cell lymphoma cells: relevance to mechanism of therapeutic action. Clinical cancer research : an official journal of the American Association for Cancer Research 8, 1234–1240 (2002).

70. Lien, E.A., Solheim, E. & Ueland, P.M. Distribution of tamoxifen and its metabolites in rat and human tissues during steady-state treatment. Cancer research 51, 4837–4844 (1991).

71. Tai, L.M. et al. Amyloid-β pathology and APOE genotype modulate retinoid X receptor agonist activity in vivo. The Journal of biological chemistry 289, 30538–30555 (2014).

72. Janowski, B.A., Willy, P.J., Devi, T.R., Falck, J.R. & Mangelsdorf, D.J. An oxysterol signalling pathway mediated by the nuclear receptor LXR alpha. Nature 383, 728–731 (1996).

73. Yang, C. et al. Sterol intermediates from cholesterol biosynthetic pathway as liver X receptor ligands. The Journal of biological chemistry 281, 27816–27826 (2006).

74. Dietschy, J.M. Central nervous system: cholesterol turnover, brain development and neurodegeneration. Biol Chem 390, 287–293 (2009).

75. Saher, G. et al. High cholesterol level is essential for myelin membrane growth. Nature neuroscience 8, 468–475 (2005).

76. Berghoff, S.A., Spieth, L. & Saher, G. Local cholesterol metabolism orchestrates remyelination. Trends in Neurosciences 45, 272–283 (2022).

77. Meffre, D. et al. Liver X receptors alpha and beta promote myelination and remyelination in the cerebellum. Proceedings of the National Academy of Sciences of the United States of America 112, 7587–7592 (2015).

